# Conservative evolution of genetic and genomic features in *Caenorhabditis becei*, an experimentally tractable gonochoristic worm

**DOI:** 10.1101/2025.05.09.653148

**Authors:** Jose Salome Correa, Luke M. Noble, Solomon A. Sloat, Tuc H. M. Nguyen, Matthew V. Rockman

## Abstract

*Caenorhabditis* nematodes represent a promising model clade for evolutionary genetics and genomics, but research has focused on the three androdioecious species, those with self-fertile hermaphrodites, all in the Elegans Group of species. The majority of *Caenorhabditis* species are gonochorists, with males and females, characterized by inconveniently high heterozygosity and inbreeding depression. We have identified *C. becei* as an experimentally tractable gonochorist. We show that transmission-genetic properties known from the androdioecious Elegans Group species are conserved in this Japonica Group species: the organization of the genetic map, cosegregation of autosomal indels and sex chromosomes, and segregation distortion due to *Medea* elements, demonstrated here for the first time in a gonochoristic *Caenorhabditis* species. We describe a new chromosomal genome assembly of a healthy inbred *C. becei* reference strain, integrating data from PacBio HiFi reads, Illumina short reads, genetic linkage, and HiC chromatin contacts, and experimental gene annotation with short- and long-read data. Some aspects of the genome are highly conserved, including synteny across the six chromosomes and the distributions of repetitive sequences and genes along each chromosome. Other features are quite distinctive, including evolved shifts in GC composition & heterogeneity along the genome. Both codon & amino acid usage are shifted in concert with the species’ genomic GC content. *C. becei* has an unusually large X chromosome, which we find is associated with multiple local gene family expansions. These findings and resources lay the foundation for further experimental and computational studies of *Caenorhabditis* genetics and genomics.

**Significance Statement:** No animal is better understood at the cellular and molecular level than the microscopic nematode *C. elegans*. To turn this knowledge to evolutionary questions, we have domesticated *C. becei*, a related species that differs in mating system. This paper describes the genomic and genetic features of this *C. elegans* relative and applies these observations to questions about evolution.

## INTRODUCTION

*Caenorhabditis elegans* is among our most powerful experimental model organisms, and decades of intensive study have revealed innumerable details about its cellular, molecular, and developmental biology. *C. elegans* sits within a speciose clade of morphologically similar nematodes that share many features, down to the number and arrangement of cells (Memar *et al.,* 2019; Zhao *et al.,* 2008; Large et al. 2025; Toker et al. 2025). Some pairs of reproductively isolated species completely lack distinguishing morphological features (Sudhaus & Kiontke, 2007). Most *Caenorhabditis* species share ecological attributes as well, reproducing within patches of decomposing plant matter and feeding on bacteria. Despite the morphological and ecological conservatism of *Caenorhabditis* nematodes, they are extremely divergent at the sequence level; distantly related *Caenorhabditis* are more diverged at the level of protein sequences than the most distantly related vertebrates or insects (Kiontke *et al.,* 2004).

The combination of conservation and divergence makes *Caenorhabditis* a particularly attractive model for comparative genomics and genetics, as factors that can confound evolutionary analysis in other groups – life history, trophic level, body size, and so forth – hardly vary within this clade (Kiontke *et al.,* 2011; Stevens *et al.,* 2019). Moreover, comparative experimental analysis of many species under identical conditions is straightforward, as more than 80 species are available in culture and compatible with standard laboratory growth conditions.

Despite these advantages, comparative experimental, quantitative, and population genetic studies have focused largely on a handful of species closely related to *C. elegans*, within the Elegans Group in the phylogeny. Further, the bulk of research has focused on the three Elegans-Group species, *C. elegans, C. briggsae,* and *C. tropicalis,* that exhibit androdioecy, a mating system of self-fertile hermaphrodites and cross-fertile males.

To understand which genetic and genomic features of the *Caenorhabditis* model species are shared more broadly across the clade, and to establish an obligately outcrossing worm as a research model, we generated a contiguous genome assembly and characterized transmission-genetic properties in *C. becei*, an experimentally advantageous species in the Japonica group, sister to the well-studied Elegans group (Stevens *et al*., 2019).

*C. becei* has many useful properties as an experimental representative of both the Japonica Group and gonochoristic (male/female) *Caenorhabditis* species more broadly. The most recent common ancestor of the Japonica and Elegans Groups (and hence of *C. becei* and *C. elegans*) is estimated to have lived approximately 720 million generations ago, versus about 530 million generations for the most common recent ancestor of *C. elegans* and either of its usual comparators, *C. briggsae* and *C. tropicalis* (Fusca *et al*. 2025). *C. becei* shares with *C. elegans* (and most *Caenorhabditis*) a compact genome, free of repetitive centromeres or Y-chromosomes, and it is amenable to cryopreservation. It grows well in laboratory culture and so has already proved useful for experimental studies of viral susceptibility, sex ratio, and reproductive biology (Huang *et al*., 2023; Lamelza *et al*., 2019; Shaw & Kennedy, 2022). Its genes and genome can be manipulated by microinjection of plasmids or nucleoprotein particles, and it has a rapid generation time, cycling from egg to egg in less than 48 hours at its native temperature of 25°C.

Phylogenetically, the species falls within a clade of *Caenorhabditis* species endemic to the Neotropics, each with a narrow geographic range (Rockman *et al*. 2026). *C. becei* is named for its type locality, Barro Colorado Island (BCI), Panamá (Stevens *et al*., 2019). It is a common species in the rainforest of BCI, the most intensively studied tropical forest on earth, and consequently it holds great promise for ecological studies (Sloat *et al*., 2022).

*C. becei*’s greatest virtue, however, and the one that motivated its selection as our model species, is that its populations harbor a relatively modest load of deleterious recessive variants, facilitating construction of healthy and fecund inbred lines (Rockman *et al.,* 2025). For short-lived small-bodied organisms like nematodes, inbred lines are a critical tool for genetic analysis, allowing experimental replication of genotypes – including outbred genotypes generated by crossing inbred lines (Andersen & Rockman, 2022). *C. becei* provides a route to genomic analysis of inbreeding depression, a phenomenon absent from the self-fertile species like *C. elegans* and nearly intractable in hyperdiverse Elegans Group species like *C. brenneri* and *C. remanei* (Adams *et al*., 2022; Baer *et al.,* 2010; Barrière & Félix, 2005; Dolgin *et al*., 2007), and to quantitative genetic analysis of outcrossing-related traits, including reproductive behaviors that are vestigial in *C. elegans*.

Previously, we reported a short-read draft assembly of *C. becei* inbred line QG2083 in 1,567 scaffolds (Stevens *et al*., 2019), and we subsequently used linkage data to consolidate the reference into a 256-scaffold assembly (Lamelza *et al*., 2019). These fragmented assemblies have already proved valuable in comparative studies of gene and regulatory network evolution (Carelli *et a*l., 2022; Eurmsirilerd & Maduro, 2020; Kursel *et al*., 2021; Maduro, 2020; Nelson & Ambros, 2021; Fusca *et al*. 2025; Jhaveri *et al*. 2025; Aharonoff *et al*. 2025). Here, we generate a new assembly, with chromosomal scaffolds, for inbred line, QG2082. We integrated data from PacBio HiFi reads, Illumina short reads, genetic linkage, and HiC chromatin contacts.

Taking advantage of *C. becei*’s compatibility with laboratory conditions, we investigated whether several transmission-genetic properties that are well characterized in the Elegans Group are conserved with this Japonica Group species. These include the organization of the genetic map into discrete recombination rate domains and the presence of segregation distortion due to *Medea* elements. These are gene-drive elements that are common in the selfing *Caenorhabditis* species but not known previously from gonochoristic species. We also tested for conservation of one of the most idiosyncratic aspects of *C. elegans* transmission genetics, the cosegregation of autosomal indel polymorphisms and sex chromosomes during male meiosis. This feature of meiosis causes autosomal indels to exhibit a sort of genetic linkage with the X chromosome, a deviation from the Mendelian expectation that separate chromosomes assort independently.

Finally, we describe the experimental and computational annotation and evolutionary analysis of genes, gene families, and repetitive elements across the *C. becei* genome, integrating these features with our newly characterized genetic map data and comparing them with results from related species. The genome of *C. becei* is highly syntenic with other *Caenorhabditis* species, but its genome exhibits several distinctive features, including high and highly variable GC content that is structured along the recombination domains, and an expansion of the X chromosome driven by the growth of gene families specifically in the chromosome arm domains. These results lay the foundation for experimental, computational, and ecological studies of genetics and genomics in *Caenorhabitis becei* and gonochoristic *Caenorhabditis* more generally, while also introducing a new arena for studies of genome evolution with relevance beyond our favorite model worms.

## RESULTS

### The genome of *C. becei*

We assembled a chromosomally scaffolded reference genome for *C. becei* strain QG2082. This inbred line was derived by 25 generations of sib-mating from wild-caught isofemale line QG704, which had been isolated in 2012 from a *Gustavia superba* flower rotting on the forest floor on Barro Colorado Island and cryopreserved within a few generations (Sloat *et al.,* 2022). Assembly integrated PacBio HiFi long reads, Illumina short reads, linkage data from a genetic cross, and HiC chromatin contact data.

The nuclear genome assembly spans 93.9 Mb. DNA sequence statistics, such as GC content, repeat content, and gene density, show symmetrical patterns along the chromosomes and align with recombination rate domains, consistent with good completeness and an absence of large-scale assembly errors. Most chromosomes terminate at both ends in oriented telomeric repeat sequences (Figure S1). Chromosomes were assigned identities based on conserved synteny with *C. elegans*. The X chromosome is much larger than the other chromosomes, almost twice the length of the shortest autosome, and 40% longer than the *C. elegans* X (Figure 1).

**Figure 1.**
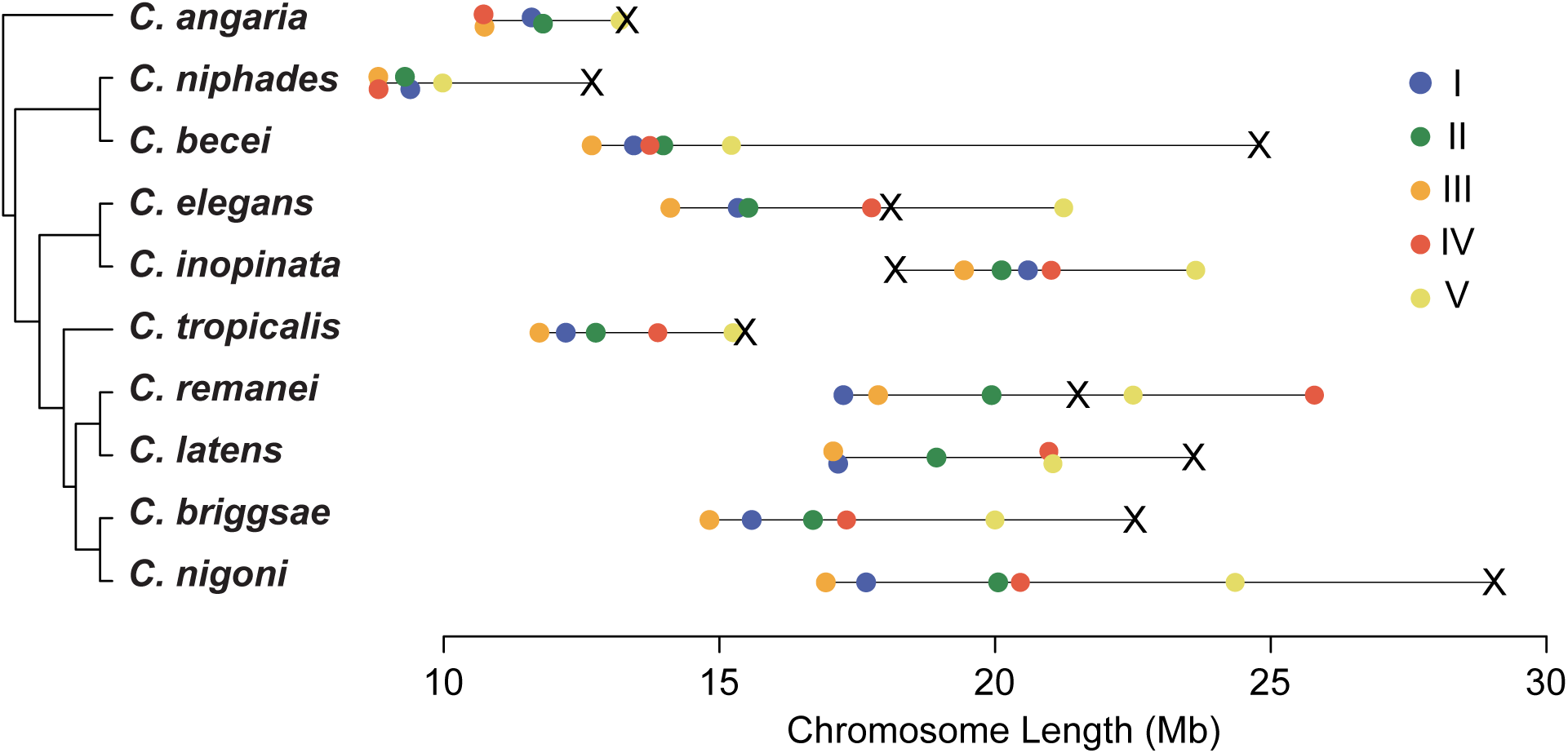
*C. becei* has a small genome but a giant X chromosome. Chromosome sizes are plotted for species of *Caenorhabditis* with published chromosomally scaffolded genomes. Here *C. niphades* and *C. becei* are the representatives of the Japonica Group; its sister clade is the Elegans Group.

Using evidence from short- and long-read transcriptome sequencing, we annotated 24,025 protein-coding transcripts representing 22,965 unique genes (File S1). Coding sequence spans 32.8% of the genome.

The genome has a low repetitive element content (15%), among the lowest in *Caenorhabditis*, comparable to *C. niphades* (11.3%), *C. bovis* (13%), and *C. elegans* (18%) (Sun *et al*., 2022; Woodruff & Teterina, 2020). The most abundant repeats were Tc1-Mariners (6.6% of the genome), consistent with their dominance across other *Caenorhabditis* species (Woodruff & Teterina, 2020). Other abundant repeat types included CACTA elements (1.3%), simple repeats (1.1%), and unclassified repeats (4.2%) (Figure S2).

To estimate repetitive element age, we used Kimura distances as a proxy for divergence. Repeat divergence distributions showed no peaks indicative of bursts of transposon expansion, and we found no evidence of recent repetitive element activity for any repeat type.

The mitochondrial genome of *C. becei* is a 13,708 bp circular molecule encoding 12 protein-coding genes, 2 ribosomal RNAs, and 22 transfer RNAs. The genome shares many features with *C. elegans*, with identical gene order and minimal intergenic space.

### Conserved architecture of the genetic map

We characterized the relationship between the genetic map and physical genome, using data from an advanced intercross between inbred lines QG2082 and QG2083 (Lamelza *et al.,* 2019). These inbred lines were derived from independent isofemale lines collected from localities separated by 3 kilometers. The resulting *C. becei* recombination landscape, the first for a *Caenorhabditis* species outside the Elegans group, closely matches the patterns known from *C. elegans*. Each chromosome has a low-recombination central domain flanked by high recombination arms and, in most cases, small tip regions with little or no recombination (Figure 2A, Table S1).

**Figure 2.**
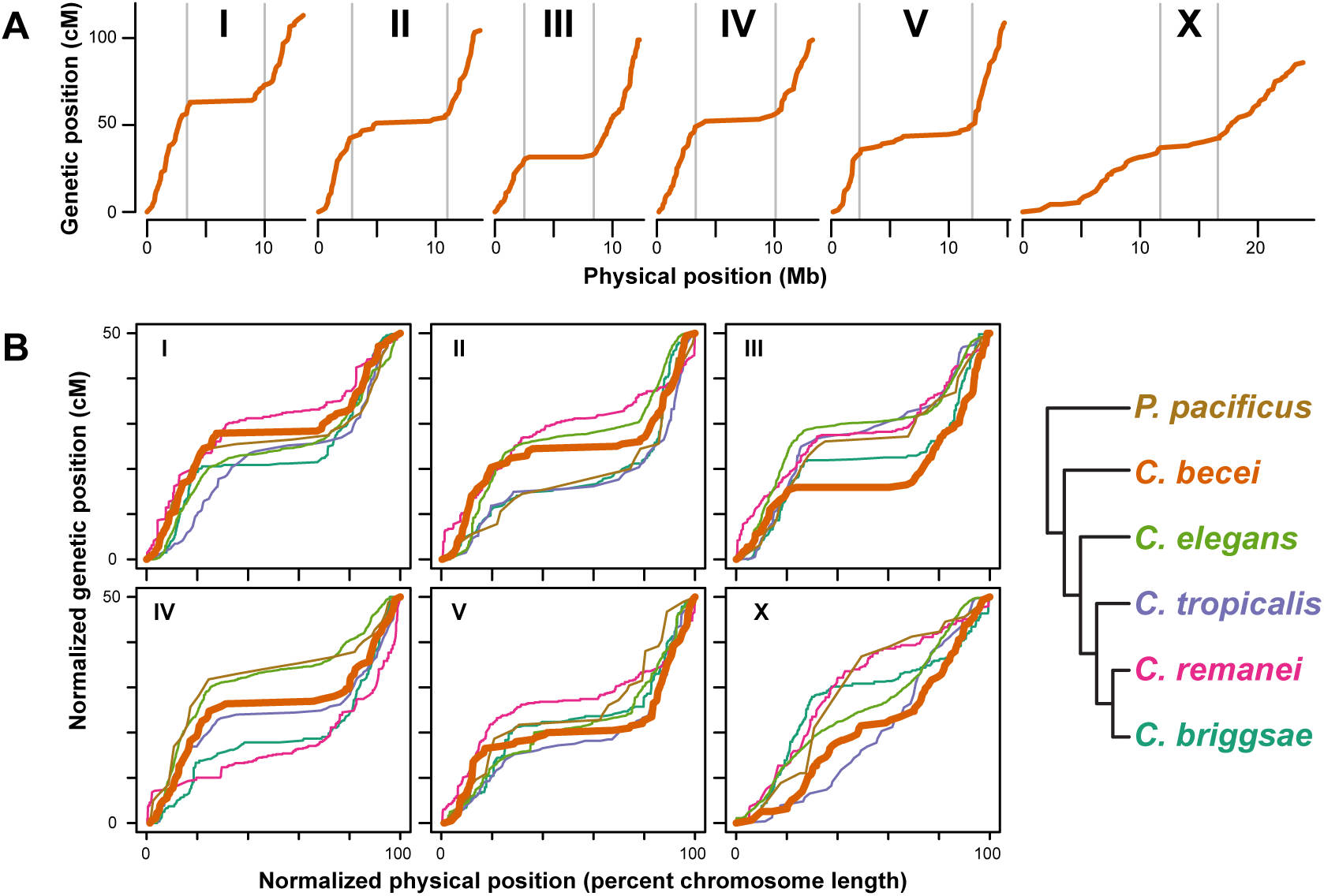
**A.** *C. becei* Marey maps showing effective genetic position (crossovers per chromosome in the G_4_BC_2_ mapping cross) as a function of physical position. Gray bars indicate the arm-center boundaries as estimated by segmented linear regression. **B.** Normalized Marey maps for the six Rhabditid species with complete physical and genetic maps. The phylogeny at right is not to scale; *Pristionchus pacificus* is very distantly related to *Caenorhabditis*.

The overall pattern of recombination rate variation along the autosomes is largely conserved across the *Caenorhabditis* species with characterized recombination maps (Figure 2B). We used segmented linear regression to estimate boundaries between the center and arm domains and to estimate recombination rates for each species in a uniform way (Table S2, Figure S3). *C. elegans, C. briggsae,* and *C. tropicalis* show concordant domains of zero, high, and low recombination in tips, arms, and centers respectively (Barnes *et al*., 1995; Noble *et al*., 2021; Rockman & Kruglyak, 2009; Ross *et al*., 2011; Stevens *et al*., 2022); *C. remanei* is exceptional; it shows the typical arm-center organization but has high recombination in the tips (Teterina *et al.,* 2023). Distantly related *Pristionchus pacificus* also shows *Caenorhabditis*-like domain architecture, despite substantial differences in meiotic machinery (Rillo-Bohn *et al*., 2021; Yoshida *et al.,* 2023).

*C. becei* resembles *C. elegans* and *C. briggsae* in having a distinct if subtle domain structure on the X chromosome, while other species have idiosyncratic or inverted X chromosome domain structure (though the *P. pacificus* X chromosome map may be distorted by inversions: Rillo-Bohn *et al*., 2021).

By estimating recombination rates for each domain (assuming 50cM chromosomes; Table S2), we found that *C. becei* autosome arms have relatively high recombination rates (mean 8.1 cM/Mb, range 6.5-10.9), similar to those of *C. tropicalis* (mean 8.2 cM/Mb) and *C. briggsae* (mean 7.8), slightly higher than those of *C. elegans* (mean 6.5) and much greater than found in *C. remanei* (mean 3.3) (Noble *et al*., 2021; Parée *et al.,* 2025; Rockman & Kruglyak, 2009; Ross *et al*., 2011; Snoek *et al.,* 2019; Teterina *et al*., 2023). These rates reflect the relative physical lengths of the chromosomes: shorter chromosomes typically have more recombination per physical length. In a naïve regression of arm recombination rate on chromosome length, incorporating all autosome arms from the five *Caenorhabditis* species, chromosome length explains 35% of the recombination rate variance (*P*<10^−5^). The relationship between rate and length is also influenced by the proportion of chromosome with crossover suppression in the center, which varies among species and chromosomes. *C. becei* autosomes have generally typical center sizes, representing approximately half the chromosome length, with the exception of the unusually long center of chromosome V (63%). However, the degree of crossover suppression in *C. becei* is unusually strong, with estimated recombination rates of only 0.1 to 0.6 cM/Mb, lower than that observed in other species (Table S2).

### Cosegregation of indels and sex chromosomes is conserved in *C. becei*

*Caenorhabditis* nematodes exhibit a remarkable pattern of correlated segregation of sex chromosomes and autosomal insertion-deletion polymorphisms (T. S. Le *et al*., 2017; Wang *et al.,* 2010). Males in these species have an X0 sex chromosome composition. In male meiosis, heterozygous autosomal insertion alleles tend to segregate away from the X chromosome and into X-nullosomic sperm, while the deletion allele segregates with the X. This pattern, known as skew, has been observed in several Elegans-group species and one more distantly related species, *C. portoensis*, but has not previously been examined in the Japonica group.

We generated an autosomally integrated transgene, *qgIs7*, which drives strong expression of fluorescent reporters. We tested whether *qgIs7* segregated at equal frequencies into male and female progeny of an F_1_ male crossed to a wild-type female. The genetic background of all strains is QG2082, and strains are expected to differ only in the presence or absence of the multicopy transgene array on an autosome. Under the skew model, the transgene should disproportionately segregate into sperm that lack an X chromosome, and therefore it will occur in an excess of male offspring and a deficit of female offspring. We observed this pattern (1,719 offspring: 343 *qgIs7* males, 278 wt males, 507 *qgIs7* females, and 591 wt females; Fisher’s Exact Test *p* = 0.0004). The Transmission Bias Ratio (Le *et al*., 2017), the ratio of preferred to unpreferred segregation, is 1.19, similar to values observed in other gonochoristic *Caenorhabditis* and much lower than in the selfing species. Another way to state these results is that the autosomal insertion is genetically linked to the nullosome at a distance of 45.7±2.4 cM.

### *C. becei* exhibits segregation distortion due to *Medea* elements

The genetic map revealed strong allele frequency skew on chromosomes I and III, with the QG2083 allele having increased in frequency in both cases (Figure 3A). The simplest model for this pattern is selection for alleles that increase developmental rate or brood size. However, QG2082 exhibits both faster development and larger brood sizes than QG2083, implying that selection favoring QG2083 alleles may have a more complicated basis. In all three selfing *Caenorhabditis* species, transmission-ratio distortion observed in experimental mapping crosses is attributable to parent-by-offspring genetic interactions, where a heterozygous parent (mother for *Medea* elements, father for *peel* elements) damages offspring that are homozygous for the non-*Medea/peel* allele (Ben-David *et al*., 2017, 2021; Noble *et al*., 2021; Rockman, 2025; Seidel *et al*., 2008; Zdraljevic *et al*., 2024). The names *Medea* and *peel* are applied to any loci with these inheritance patterns and do not imply homologous mechanisms or shared ancestry.

**Figure 3.**
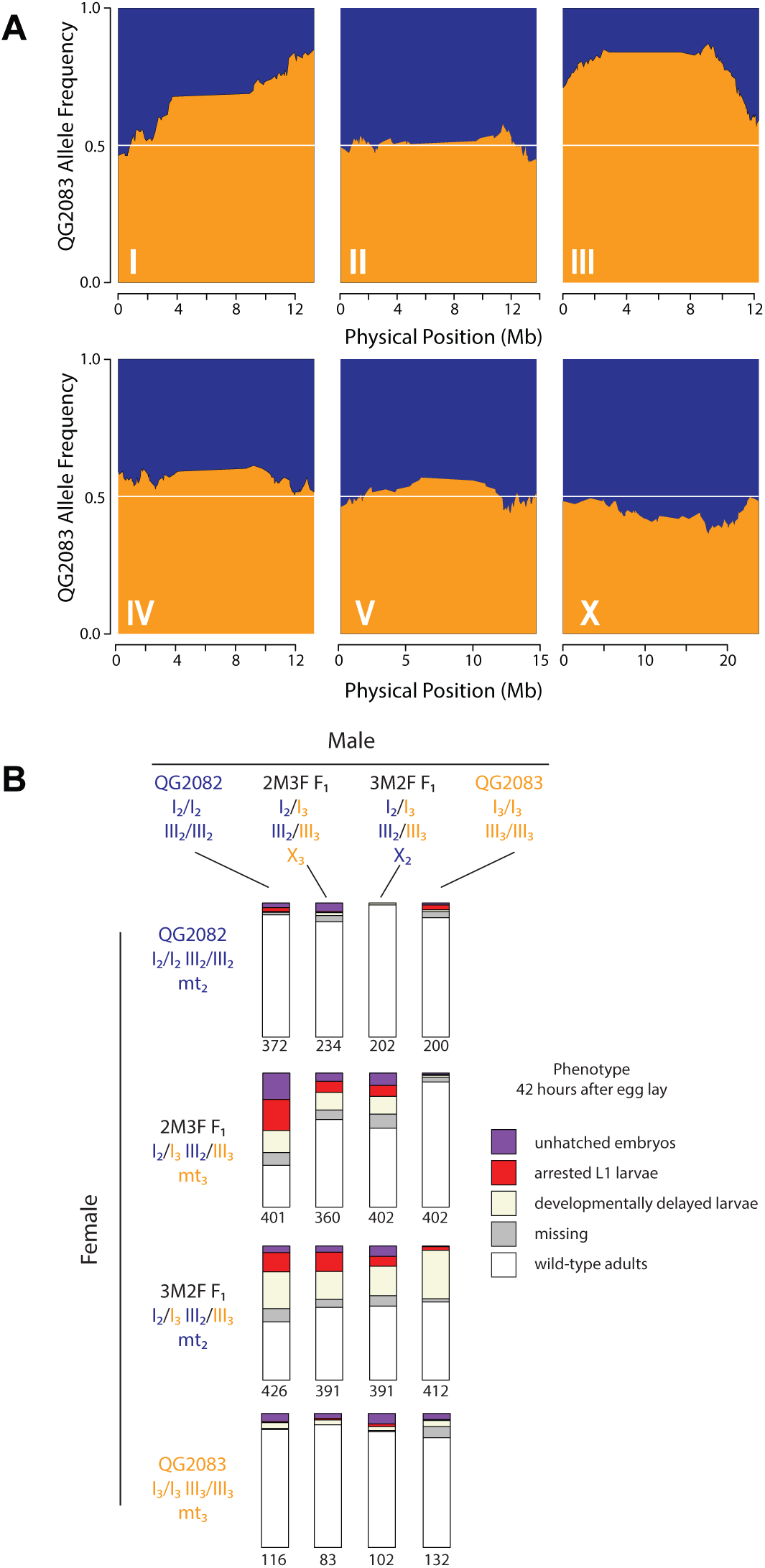
*Medea* alleles cause segregation distortion. **A.** Allele frequencies among G_4_BC_2_ recombinant genomes. Strong distortion of inheritance favoring QG2083 is present on chromosomes I and III. **B.** Experimental crosses implicate maternal-effect loci that act in heterozygous mothers and cause lethality or arrest in homozygous offspring. The phenotype frequencies of progeny from each of 16 classes of cross are shown in stacked bar charts, with sample size below. Parental genotypes on chromosomes I, III, X, and mitochondrial genomes are colored according to their origins in QG2082 or QG2083, and also indicated by the subscripts 2 or 3, respectively.

We performed experimental crosses to test for *Medea* or *peel* activity, and we discovered that QG2083 carries alleles with *Medea* activity (Figure 3B). Only offspring of heterozygous mothers were affected, and the affected offspring were in proportion to the expected fraction homozygous for QG2082 alleles (Figure S4). Offspring of F_1_ females x QG2082 males were most severely affected, with less than half of progeny showing wild-type development, while the reciprocal crosses, QG2082 females x F_1_ males, showed no impact. By comparing the counts of affected offspring from different classes of crosses, and assuming that the loci on chromosomes I and III are each acting as *Medea* loci, we estimate that each has a penetrance exceeding 50%. The data also show a potential interaction with mitochondrial genotype or a parent-of-origin effect, as also observed for *Medea* elements in *C. tropicalis* (Ben-David *et al.,* 2017, 2021; Noble *et al*., 2021), seen in the different patterns in the second and third rows of Figure 3B.

### Nonuniform distribution of genes and repeats

Having characterized the evolutionary conservation of transmission genetic properties of *C. becei* above, we now turn to questions about conservation of genome sequence features. In *C. elegans* and other well studies species, genome annotations are highly structured according to chromosomal domain, and *C. becei* follows these patterns. Repetitive elements are more abundant in the arms than in the centers (Figures S5). This non-random distribution of repeats is consistent across most *Caenorhabditis* species, except for *C. inopinata* and *C. bovis*, where a more homogeneous repeat landscape is observed due to the recent expansion of Tc1-Mariner elements (Woodruff & Teterina, 2020). Gene densities are also nonuniform along the chromosomes (Figure S6). Coding sequence and non-repetitive intergenic sequences are densest in chromosome centers, while chromosome arms show higher densities of both intronic DNA and repeat sequence.

### Genomic GC Content

Among species with chromosomal assemblies, *C. becei* has genomic GC content that is both high, at 42.4%, and unusually variable along the chromosomes (Figure 4). The variance in GC content along chromosomes is much greater in *C. becei* than in the other species with chromosomal assemblies, independent of the scale over which GC% is measured (Figure S7). Chromosomes I-IV exhibit notable contrasts between high GC content in their centers and lower GC content in their arms. In contrast, chromosomes V and X show a more uniform distribution of high GC content along both the center and the arms.

**Figure 4.**
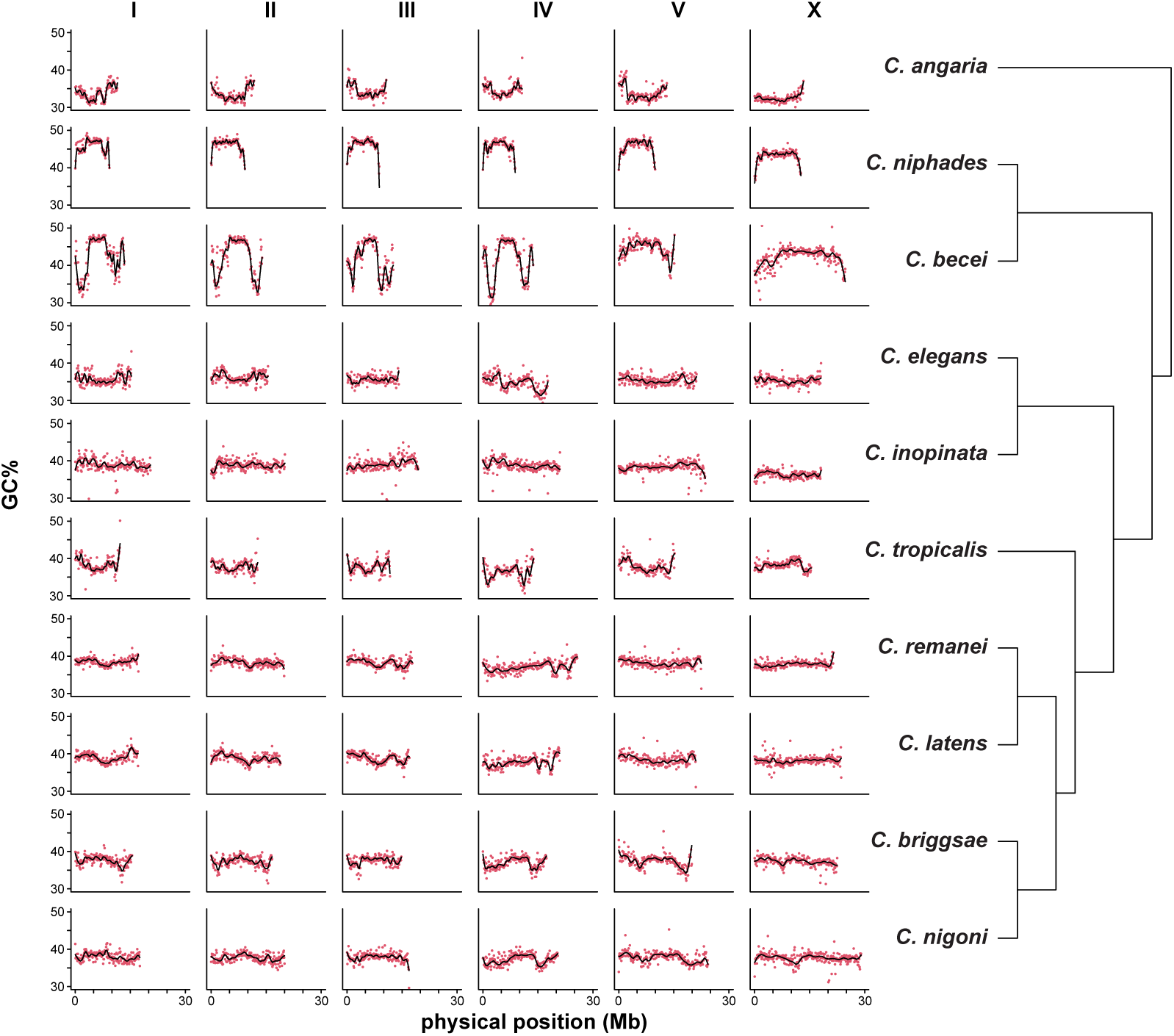
GC content varies along chromosomes and among species. GC% is plotted for non-overlapping 100kb windows along the length of each chromosome, along with LOESS-fitted lines (degree 2, span 1/8). *C. becei* is unusual in its high GC content (a trait shared with *C. niphades*, the other Japonica Group species with a chromosomal assembly) and its high variance, most notably on chromosomes I-IV.

The average GC content of features follows a descending order of exon > genomic average > repeat > introns/intergenic regions, and the pattern holds in both chromosome arms and centers (Figure S8). This ordering is conserved across species, but the differences are more pronounced in *C. becei* and Elegans Group species than in *C. niphades* or *C. angaria* (Figure S8). In these latter species, GC content is relatively consistent across the chromosomes within each species and across annotation features.

### Codon and Amino Acid Usage

Codon usage varies among *Caenorhabditis* species, tracking GC content variation. *C. becei* and *C. niphades* have high GC content and use codons with higher GC content, while *C. angaria* has lower GC content and uses codons with lower GC content (Figure 5A). Species in the Elegans group exhibit intermediate codon usage patterns.

**Figure 5.**
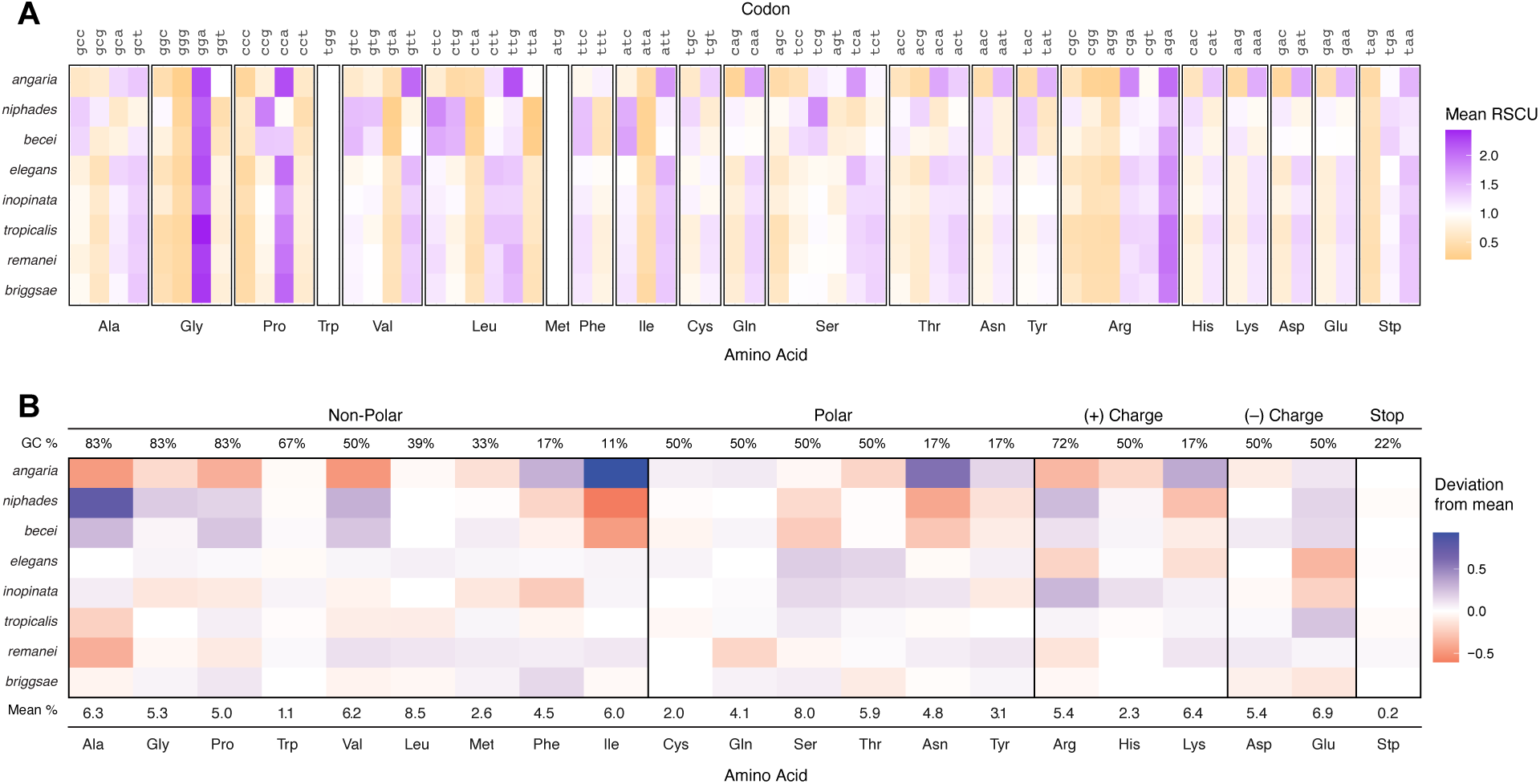
Codon usage and amino acid composition in *Caenorhabditis* species. **A.** Relative synonymous codon usage (RSCU). Rows correspond to species, and columns represent codons grouped by amino acid. Tiles represent the RSCU value for each codon in a given species. RSCU values greater than 1.0 indicate codons used more frequently than expected under uniform usage, with higher values shaded in orange and lower values shaded in purple. **B.** Amino acid usage: Rows correspond to species, with the last row displaying the mean percentage of each amino acid across all species. The mean is calculated by averaging the percentage of each amino acid in the CDS across species. Tiles represent the deviation from the mean amino acid percentage for each species, with the mean calculated separately for each amino acid. Fill colors indicate the direction and magnitude of the deviation: values above the mean are shaded in blue, and values below the mean in red. The intensity of the shading reflects the magnitude of the deviation. The top row shows the average GC% of codons encoding each amino acid.

A similar trend is observed in amino acid usage: *C. becei* and *C. niphades* show positive deviations from the mean for amino acids associated with high GC codons and negative deviations for those linked to low GC codons. In contrast, *C. angaria* exhibits the opposite pattern, with positive deviations for amino acids associated with low GC codons and negative deviations for those linked to high GC codons (Figure 5B).

### Synteny

We observe strong synteny between the chromosomes of *C. becei* and *C. elegans* (Figure 6), as expected given the well-established conservation of ancestral linkage groups (Nigon elements) in Rhabditid nematodes (Tandonnet *et al*. 2019; Gonzalez de la Rosa *et al.,* 2021). Using 9,833 1-to-1 orthologs between these species, 9,657 pairs (98.2%) lie on chromosomes with matching identities, confirming nearly complete conservation of chromosome-scale linkage. Within this overall pattern, the most striking feature is the pronounced deficit of 1-to-1 orthologs across broad regions of the *C. becei* X-chromosome arms, which coincide with domains rich in gene duplications, discussed below in the section on the distribution of orthologs and paralogs.

**Figure 6.**
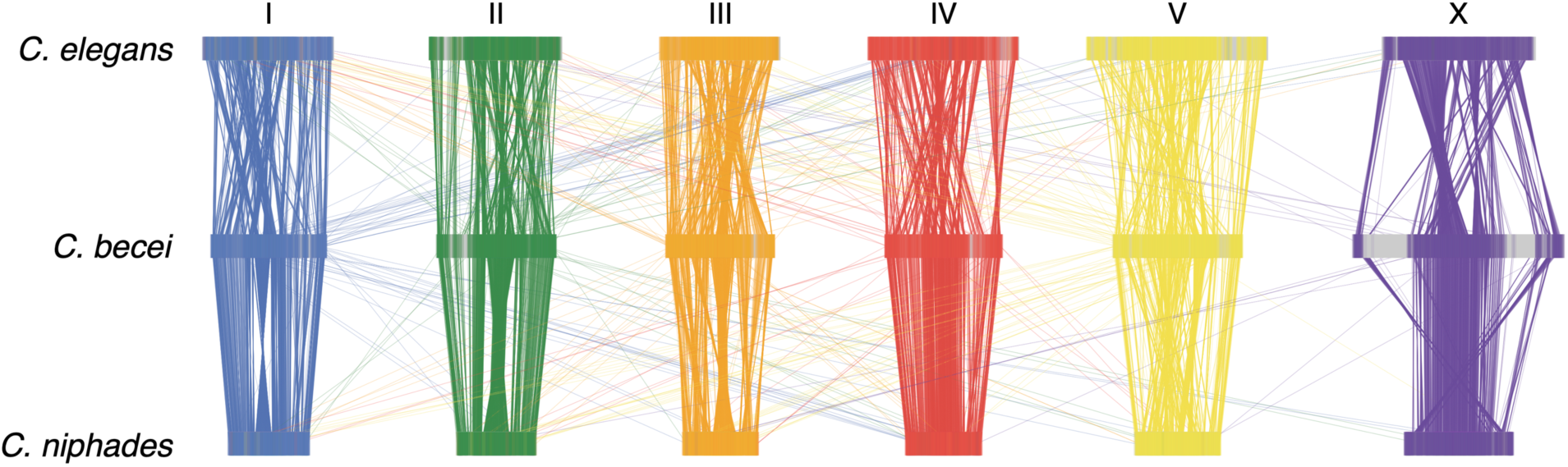
Synteny between *C. elegans*, *C. becei,* and *C. niphades*. Each line connects single-copy orthologues, colored according to *C. becei* chromosome.

We further compared synteny between *C. becei* and *C. niphades*, the closest relative with a chromosomal assembly, and between *C. elegans* and *C. niphades* (Figure 6). The *C. becei*–*C. niphades* comparison includes 9,710 1-to-1 orthologs, the *C. elegans*–*C. niphades* comparison includes 10,224 1-to-1 orthologs, and 8,777 genes form simple 1-to-1-to-1 ortholog trios shared among all three species. In each species pair, approximately 98% of 1-to-1 orthologs reside on homologous chromosomes. On Chromosome X, nearly all 1-to-1 orthologs between *C. becei* and *C. niphades* also map to the X in both genomes, but the criss-crossing links in the synteny plot indicate extensive reordering of gene positions relative to *C. elegans*. The X-linked regions where *C. becei* lacks single-copy orthologs with *C. elegans* are likewise depleted of 1-to-1 partners in *C. niphades*, so these *C. becei* X-arm domains fall outside the conserved single-copy ortholog backbone shared among the three species.

### Comparative ortholog analysis reveals widespread gene duplications in *C. becei*

We used gene annotations from high-quality chromosomal assemblies of ten Elegans Supergroup species to classify genes into orthogroups. Orthogroups were categorized as single-copy, multi-copy, or unassigned. Multi-copy orthogroups contained more than one gene within a species, while single-copy orthogroups included genes that were present as a single copy in a given species and had orthologs in other species. Unassigned orthogroups consisted of genes that were species-specific, with no detectable homologs.

Across the 22,965 protein-coding genes in *C. becei*, 11,790 (51.3%) were classified as single-copy, 9,537 (41.5%) as multicopy, and 1,638 (7.1%) were unassigned, *C. becei*-specific genes in this sample of genomes. These numbers reveal an elevation in multicopy genes in *C. becei* compared to *C. elegans* (6,716 multicopy genes, 33.6%) and *C. niphades* (3,611 multicopy genes, 22.2%). In each species, but especially in *C. becei*, multicopy genes are concentrated in chromosome arms and single-copy genes in chromosome centers and tips (Figure 7). Multicopy genes are also concentrated on chromosome V relative to the other autosomes. Finally, the *C. becei* X is exceptional for both its high proportion of multicopy genes and their concentration on the arms. More than 52% of genes on the *C. becei* X are multicopy, versus 20% in *C. elegans* and 22% in *C. niphades*. As seen in Figure 7, the large regions on the X-chromosome arms that are devoid of syntenic 1:1 orthologs are not gene deserts but rather thickets of multicopy genes.

**Figure 7.**
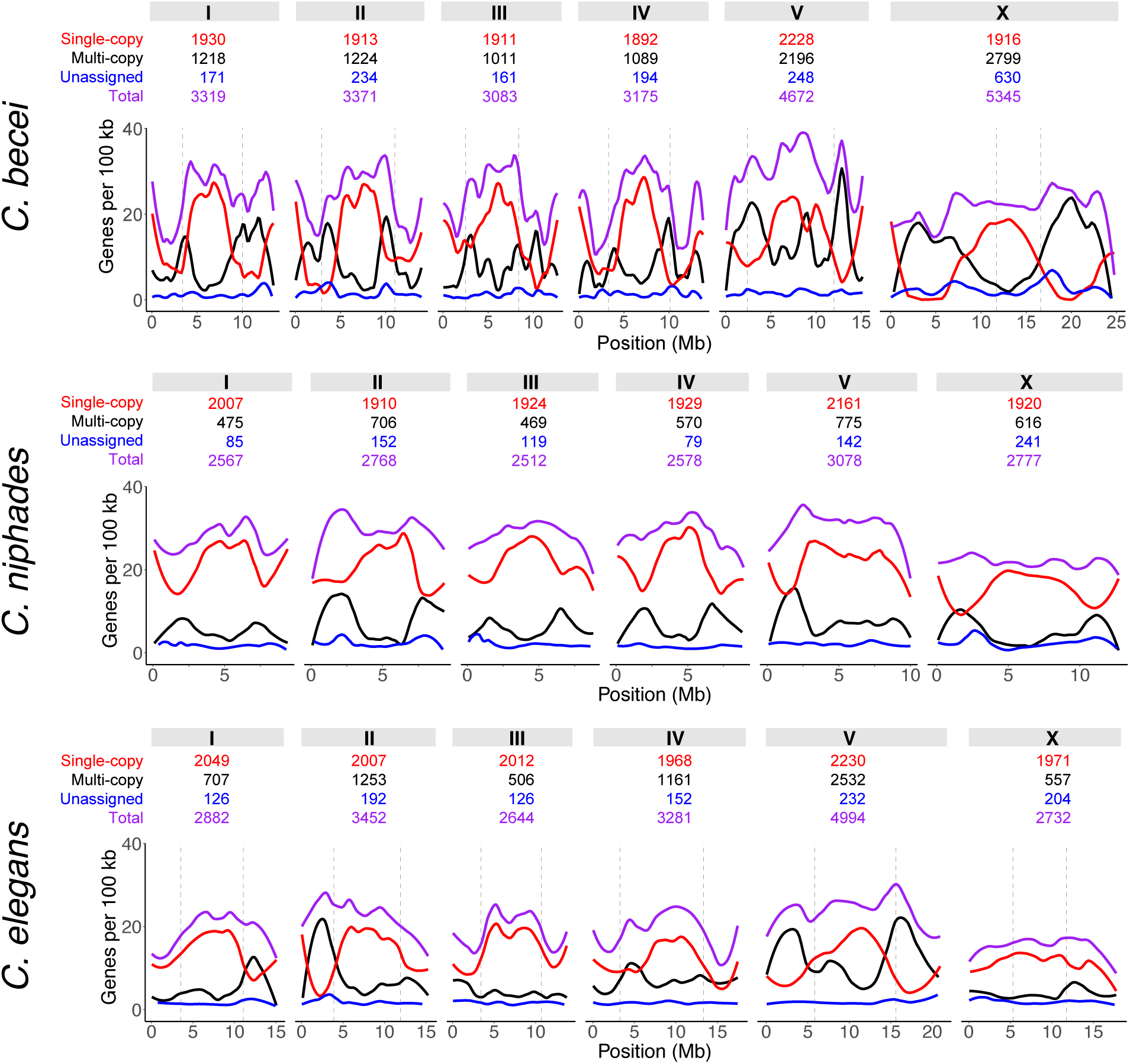
Distribution of single-copy, multicopy, and unassigned genes across the chromosomes of *C. becei*, *C. niphades*, and *C. elegans*. Gene counts are plotted in 100 kb windows across the chromosomes of each species, with LOESS-fitted lines (span = 0.2). Chromosome arm-center boundaries from recombination maps are plotted as dotted lines.

Genes within multicopy orthogroups tend to occur in local clusters (Figure S9). Among the 88% of multicopy genes that are on the same chromosome as a same-orthogroup paralog, 4,636 genes (55.5%) fall within 5kb of the nearest paralog, 2,045 (24.5%) fall between 5 and 50 kb, and 1,678 (20.1%) are separated by more than 50 kb.

We characterized the *C. becei* proteome by annotating protein domains in each of the 22,965 genes (File S2). Overall, 84.1% of genes are assigned InterPro domain annotations, ranging from 90.3% among single-copy genes to 81.3% in multicopy genes and 56.5% in unassigned (*C. becei*-specific) genes. Among the multicopy orthogroups, GPCRs and F-box domains are most abundant domains, along with DUF38/FTH domain proteins, lectins, and Nuclear Hormone Receptors (Figure S10).

Protein domains define gene families that vary strikingly in their patterns of spatial clustering (Figure S11). For example, DUF38/FTH-domain-containing genes are only 4.4% single-copy and 78.9% closely-clustered (<50 kb) multicopy genes. F-box, T-box, and MATH-domain genes show similar biases toward local multicopy gene clusters. By contrast, some domains are predominantly represented by single-copy gene orthogroups. These include C2H2-type zinc fingers (84.7% single-copy), WD40-repeat domains (87.5%), and collagens (72.4%).

The distribution of gene families is also highly structured by chromosome (Figure S12). For example, of multicopy genes that contain GPCR domains, 541 of 751 (72%) are on chromosome V. Similar concentrations of multicopy genes on V are observed for MATH-domain genes, BTB/POZ-domain genes, and NHR genes. The X chromosome carries concentrations of multicopy genes with roles in gene regulation, including T-box genes (50 of 84) and homeobox genes (45 of 80).

The evolutionarily distinctive arm regions of the X chromosome (here defined as 100 kb windows with less than 20% single-copy gene content) contain 2,049 multicopy genes, representing 639 different orthogroups, as well as 356 unassigned genes, together accounting for more than 10% of all the genes in the *C. becei* genome. Among the multicopy genes in these regions, 609 lack a protein domain or feature annotation, while intrinsically disordered regions and coiled-coil features occur in 915 and 413 genes, respectively. The most frequently represented domain families include F-box (78 genes), T-box (50), DUF38/FTH domain (44), Homeobox (41), nematode-specific protein (39), and CUT (36) domains. Because individual genes can contain more than one distinct domain family, these counts can overlap; for example, many CUT-containing genes also contain a Homeobox domain. In short, the large size of the X chromosome reflects a large number of gene-family expansions involving hundreds of different orthogroups carrying a broad range of domain families.

## DISCUSSION

*Caenorhabditis becei* is a promising model for studies of gonochoristic *Caenorhabditis*, and we here lay the genetic and genomic foundations for future studies. Genetically, we find that *C. becei* is very much a *Caenorhabditis*, with an *elegans*-like genetic map and transmission-genetic properties. Genomically, *C. becei* shows an *elegans*-like pattern of gene-dense chromosome centers and repeat-dense chromosome arms, and no evidence for recent proliferation of transposable elements. But we see distinctive features as well, including elevated GC content, a concomitant shift in codon and amino-acid usage, and also pronounced heterogeneity in GC content along some of the chromosomes. Though the chromosomes of *C. becei* are highly syntenic with other *Caenorhabditis*, the X chromosome has experienced substantial growth associated with multiple gene-family expansions specifically in the chromosome’s high-recombination arm domains. The biological significance of these gene-family expansions emerges as a major question for future studies.

Our genetic map, like those previously generated for the androdioecious *Caenorhabditis* species, was affected by strong segregation distortion on several chromosomes. We showed that the observed segregation distortion likely reflects the action of multiple *Medea* elements, which act in heterozygous mothers and cause developmental delays or lethality in offspring that do not inherit them (Beeman *et al.,* 1992). This is the first demonstration of *Medea* elements in the Japonica group and in dioecious *Caenorhabditis*, and it suggests that the striking ubiquity of these genetic elements (Ben-David *et al.,* 2017, 2021; Noble *et al.,* 2021; Seidel *et al.,* 2008; Zdraljevic *et al.,* 2024) is a broadly shared feature of *Caenorhabditis* biology, not dependent on the androdioecious mating system of the selfing species. An analytical model of *Medea* dynamics found that alleles could be maintained in partially selfing species by local positive frequency dependence, which predicts that *Medea* alleles should be fixed within local selfing populations and different between them (Rockman 2025). In obligate outcrossers, however, *Medeas* should sweep to fixation. In our study, the alternate alleles at the *Medea* loci are from isofemale lines collected within a few kilometers of one another in 2012, and additional studies will be required to determine whether they are indeed spreading through the population on Barro Colorado Island. In the meantime, experimental geneticists working in *C. becei*, and presumably all *Caenorhabditis* species, should be alert to the potential impact of *Medea* elements.

At the DNA sequence level, the *C. becei* genome is distinctive for its high GC content, a feature shared with *C. niphades* and, based on draft genome assemblies, with closely related species in the Japonica group (Stevens *et al*. 2019; Sun *et al*. 2022). The elevated GC content influences codon usage bias in *C. becei*, as demonstrated by comparative analysis across *Caenorhabditis*: species with higher genomic GC content tend to preferentially use GC-rich codons. This bias extends beyond codon usage to amino acid composition: both *C. becei* and *C. niphades* have elevated frequencies of amino acids encoded by GC-rich codons. Synonymous codon usage is under selection in *Caenorhabditis* (Cutter *et al*. 2008), but shifts in codon usage bias are generally interpreted as consequences of changes in mutational patterns rather than changes in optimal codon identity (Cutter et al. 2006; Knight et al. 2001). Our findings are consistent with this inference and with surveys across all of Nematoda (Cutter *et al*. 2006), showing largely conserved optimal codons but divergent codon usage associated with shifts in GC content. Similarly, the relationship between GC content and amino acid usage is well established (Foster 1997; Knight *et al*. 2001; Cutter *et al*. 2006), and here implies that (among other things) standard time-reversable models of molecular evolution may be misspecified for *Caenorhabditis*.

Chromosomes I-IV (but not V or X) have dramatic heterogeneity in GC content, very high in centers and tips and very low in the chromosome arms. These arm regions are also dramatically depleted of coding sequences relative to other parts of the genome, and coding sequences are substantially more GC-rich than other classes of sequence. However, the GC pattern is not purely compositional, driven by coding sequence density; intronic, intergenic, and repetitive sequences all show the same pattern of high GC in the centers and low in the arms. The extreme variation in GC content along chromosomes I-IV may instead reflect chromosome-specific mutational processes. Note that GC-biased gene conversion is not evident in *Caenorhabditis* data (Cutter & Charlesworth 2006), and moreover it would be expected to lead to higher GC in the high-recombination arms (Teterina et al. 2023), the opposite of the observed pattern.

Repeat elements are strongly enriched in chromosome arms and depleted in centers, matching recombination rate variation across the genome. The depletion of repeats in chromosome centers is likely due to strong purifying selection, which removes insertions in these functionally constrained regions (Woodruff & Teterina, 2020). This is similar to the paucity of gene duplications in centers, where functionally constrained genes dominate, and disruptive insertions are more likely to be deleterious (Thomas, 2006). The arm-rich, center-poor repeat distribution is also observed in other *Caenorhabditis* species without recent repeat activity (Woodruff & Teterina, 2020).

The *C. becei* X chromosome is exceptionally large, 40% longer than that of *C. elegans*, and it contains more genes than any other chromosome. X chromosomes are special in multiple ways, including dosage compensation, sex-biased gene content, and hemizygosity in males, which reduces recombination and exposes recessive alleles to selection. In *C. becei*, the large size of the X is associated with a large number of gene family expansions, specifically in the high-recombination arm regions. These regions carry more than 2,400 genes that are not 1:1 orthologous with genes in *C. elegans*, about three-quarters as many genes as on the whole of the smallest autosome.

Many of the gene families on the X arms are also common among multicopy genes on autosome arms, and are known as fast-evolving gene families that are also structurally variable within species (Thomas et al. 2005; Thomas 2006; Lee et al. 2021). Gene families such as GPCRs, lectins, F-box genes, and Nuclear Hormone Receptors are enriched in the hyper-divergent regions of chromosome arms in *C. elegans, C. tropicalis,* and *C. briggsae* (Lee et al. 2021; Noble et al. 2021). Gene family diversity is also elevated in the chromosome arms of more distantly related species such as *Heligmosomoides bakeri* and *Meloidogyne hapla* (Stevens *et al*. 2023; Shakya et al. 2025). The gene families in these hyperdiverse arm regions are hypothesized to play key roles in environmental sensing and pathogen avoidance and resistance, and their diversity inferred to represent the effects of long-term balancing selection (Moya et al. 2025).

While the thousands of multicopy genes on the X likely have many stories to tell, some involving familiar classes of divergent genes functioning in environmental sensing and immunity, one unusual gene-family expansion on the X involves genes with known dosage-sensitive roles, where the number of gene copies is expected to impact phenotypes, independent of divergence among them. The X-chromosome arms contain 36 genes with DNA-binding CUT domains, versus seven in the whole *C. elegans* genome (Reilly et al. 2020); *C. becei* carries an additional 27 CUT-domain genes on its autosomes. Twenty-five of the X-arm genes are in the ONECUT subfamily, carrying a single CUT domain followed by a Homeodomain. In *C. elegans*, the six ONECUT transcription factors play two independent dosage-sensitive roles, in neuronal identity and sex determination.

First, the ONECUT genes regulate the battery of genes that are expressed in all neurons in *C. elegans*. The genes act interchangeably and cumulatively to drive expression of these pan-neuronal genes; individual mutants show similar dosage-sensitive effects, and overexpression of single genes rescues a sextuple mutant (Levya-Diaz & Hobert 2022).

Second, a ONECUT gene in an arm of the *C. elegans* X chromosome, *ceh-39*, plays an additional role earlier in development, helping cells to count their X chromosomes and activate different sex determination pathways in XX and X0 individuals (Gladden and Meyer 2007). *C. elegans ceh-39* and a few other genes on the X chromosome act as XSEs, X-Signal Elements, competing with a handful of autosomal genes, ASEs, to influence transcription and post-transcriptional regulation of *xol-1* (Farboud et al. 2013). XOL-1 activity is required for male differentiation and male-specific suppression of dosage compensation, which reduces expression in XX individuals in *Caenorhabditis.* High doses of XSEs (including CEH-39) in XX animals repress *xol-1* transcription, while the lower dose in X0 animals is overwhelmed by the greater autosomal activity of *xol-1*-activating ASEs (Meyer 2022). The correspondence between *ceh-39* and the many ONECUT genes on the *C. becei* X is unclear, but remarkably, *xol-1* is itself also expanded in *C. becei*: the gene is present in three copies on the right arm of the X chromosome.

Turnover of chromosome-counting mechanisms in *Caenorhabditis* would not be too surprising, as the precise X:Autosome threshold for sex-determination is known to differ between *C. elegans* and *C. briggsae* (Harbin *et al*. 2026), and differences between *C. briggsae* and its sister species *C. nigoni* in *xol-1* regulation, and consequently in sex-chromosome dosage compensation, are implicated in their reproductive isolation (Li *et al*. 2025; Harbin *et al*. 2026). While we speculate that the genetic architecture of chromosome counting may have distinctive properties in *C. becei*, we note that the downstream mechanisms of X dosage compensation appear to be conserved (Aharonoff *et al*. 2025), based on analysis of the HiC data we report here, implying a pattern of developmental systems drift: the molecular details evolve while their output remains constant (True and Haag 2001).

The *C. becei* genome represents an appealing model for studies of genome evolution, with interactions between gene duplication, GC content heterogeneity, recombination rate variation, and codon usage biases. Comparative analyses with additional *Japonica* species will provide further insight into which of these patterns are unique to *C. becei* and which reflect broader evolutionary trends within the clade.

## METHODS

### Genome Assembly

QG2082 was grown on NGMA plates at 25°. A mixed-stage mixed-sex population was flash frozen in liquid nitrogen and the frozen worm pellet was shipped to the University of Oregon Genome and Cell Characterization Core Facility for HiFi sequencing. Flash-frozen worms were also shipped to Dovetail Genomics (Santa Cruz, CA; https://dovetailgenomics.com) for Hi-C sequencing with 150bp Illumina reads. Short-read data for QG2082 (Illumina paired-end), QG2083 (Illumina paired-end and mate-paired), and for G_4_BC_2_ lines (Illumina paired-end) are described in Lamelza *et al*. (2019).

We generated assemblies using PacBio HiFi reads (≥Q20) with assemblers Flye (Kolmogorov *et al*., 2019), Hifiasm (Cheng *et al*., 2021), and Hi-Canu (Nurk *et al*., 2020), and Hi-C-scaffolded Flye and Hifiasm assemblies with Juicer (Durand *et al*., 2016). As in Noble *et al*. (2021), we then evaluated each assembly for consistency with genetic linkage data (implemented in a Snakemake pipeline available at https://github.com/lukemn/becei), and then closed gaps of estimated 0cM genetic distance where strand-consistent spanning reads from local mapping were available. The final assembly drew partially redundant sequences from two primary assemblies (Flye -m 10 -g 100Mb; Hifiasm -l 0). For chromosomes I, IV and X, sequences from Hi-C scaffolding of the Hifiasm assembly were used to span primary sequences. Each chromosome was iteratively polished with HiFi reads, using calls from DeepVariant (Poplin *et al*., 2018) and bcftools norm/consensus (Danacek *et al*., 2021), until either no homozygous variants were called, or a local minimum in read mapping error rate was found. Telomeric sequences were counted using R library *seqinr* (Charif & Lobry 2007).

To assemble the *C. becei* mitochondrial genome, we first mapped Q20- and Q30-filtered PacBio HiFi reads to the *C. becei* nuclear reference genome using minimap2 (version 2.22; Li 2018), and extracted unmapped reads using samtools (version 1.14; Danecek *et al.,* 2021). We then aligned unmapped Q20 reads to the mitochondrial genome of *C. nouraguensis* (Accession: NC_035250.1), using minimap2 to identify mitochondrial reads. Subsequently, we *de novo* assembled the Q20 mitochondrial reads using Flye (version 2.9.5; Kolmogorov *et al.,* 2019) with a minimum overlap of 1,000 bp, and generated a single circular 13,708 bp contig with high coverage. This draft contig was then used as a reference to map the higher-accuracy Q30 HiFi reads, which we assembled using Flye with a 10 kb minimum overlap, producing a single high-confidence circular contig. This final assembly was annotated using MITOS2 (Al Arab *et al.,* 2017; Donath et al. 2019) and manually curated.

The nuclear and mitochondrial genome assemblies and underlying data are available from the NCBI at BioProject PRJNA989223.

### Genetic Map

Data for genetic map construction are described in Lamelza *et al*. (2019). For the map described here, we used short-read sequence data from 92 individual G_4_BC_2_ populations. Each such population is a pool of worms carrying one unique set of recombinant chromosomes, each the product of several generations of meiosis. Reads were mapped to the QG2082 reference genome and ancestries inferred along each chromosome using MSG (Andolfatto *et al.,* 2011). The data plotted in Figure 3A are in Table S1.

Domain boundary positions for *C. becei* and other species were estimated by segmented linear regression using the *segmented* package (Muggeo, 2008). We first interpolated genetic positions for 200 evenly spaced physical positions along each chromosome, then excluded 1 Mb from each end of each chromosome to avoid effects of tip regions, which generally have suppressed recombination and whose boundaries are poorly resolved in most species given the limited available data. We then regressed genetic position on physical position with the constraint that there be three linear segments. Note that domain boundary positions are imprecise, given the modest number of crossovers present in the data; as shown by simulations in (Rockman & Kruglyak, 2009), even in the case of precise boundaries, uniform domains, and relatively large samples of crossovers, the confidence intervals for boundaries are on the order of a megabase.

For comparisons with other species (Table S2 and Figure S3), we used the following datasets:

|  |  |  |  |
| --- | --- | --- | --- |
| *C. briggsae* | 99 F<sub>2</sub> RILs | QX1410 x VX34 | Stevens *et al.*, 2022 |
| https://github.com/AndersenLab/briggsae_reference_genome_MS/blob/main/3_recombinationmap/CB_genetic_map.Rda |  |  |  |
| *C. tropicalis* | 119 F<sub>2</sub> RILs | NIC58xJU1373 | Noble *et al.*, 2021 |

https://github.com/lukemn/tropicalis/blob/master/geneticMap/data/c_tropicalis_flye_geneticMap.rda
*C. remanei* 341 F<sub>2</sub>s PX506 x PX553 Teterina *et al.*, 2023
https://figshare.com/ndownloader/files/41808561
*C. elegans* 1045 RIAILs CB4856 x N2 & CB4856 x QX1430 (N2 *peel-1(ttTi12715)*
I *npr-1(qg1)* X) Brady *et al.*, 2019
https://github.com/AndersenLab/linkagemapping
The X-chromosome map was re-estimated after excluding markers surrounding the *npr-1* introgression *qg1* in RIAIL sets 2 and 3, which cause a spurious map expansion otherwise.
*Pristionchus pacificus* 93 F<sub>2</sub>s PS312 x PS1843 Rillo-Bohn *et al.*, 2021

We used the *P. pacificus* oocyte genetic map data only, due to strain-dependent crossover suppression in the male data, likely attributable to inversions. The X data are also distorted in a putatively strain-dependent fashion, as described in Rillo-Bohn *et al*. (2021).

### Autosomal Sex Linkage

To create a strain with an integrated transgene insertion, we used Gibson cloning to generate plasmids pSAS02 (*Cbr-P_plg-1_::GFPnovo2::unc54utr* with the pSM-GFPnovo2 backbone) and pSAS06 (*Cel-P_fkh-6_::mCherry::unc54utr* in the pCFJ104 backbone). The backbone plasmids were acquired from Addgene and the final plasmids were verified by Oxford nanopore whole-plasmid sequencing (Plasmidsaurus.com). We PCR-amplified the reporters (*promoter::FP::UTR*) from the plasmids and injected them together into QG2082 and recovered lines transmitting both transgenes. Several hundred L4 animals of a transmitting line were UV irradiated at 500 J/m^2^. Those animals were passaged for 5 generations before backcrossing individuals to QG2082 to screen for Mendelian segregation of the integrated array. Lines positive for integration were backcrossed an additional 12x before homozygosing the integrated array as line QG4602 (*qgIs7*). Plasmid sequences are provided in File S3.

To test for skew, we performed reciprocal crosses between QG2082 and QG4602 under standard conditions at 25°C. We then crossed F_1_ males to QG2082 females and collected and scored all progeny laid within timed windows. The female-biased sex ratios we observed are consistent with the known bias in *C. becei* (Huang *et al.,* 2023). Recombination frequency is estimated as the number of unpreferred-phase gametes (X+insertion or 0+wildtype) out of *n* = 1,719 total, and the 95% confidence interval is the normal approximation for binomial proportions.

### Segregation distortion

To test for parental-by-offspring genetic interactions (*Medea* or *peel*), we performed replicates of the sixteen possible pairwise crosses between four genotypes: QG2082, QG2083, and the F_1_s from each direction of the cross between QG2082 and QG2083. We included all possible F_1_s in order to account for possible effects of mt-DNA or X-linked interactors. The reciprocal F_1_ females differ in their mitochondrial genotype, and the reciprocal F_1_ males differ in which X chromosome they carry. Under a *Medea* hypothesis, the only affected individuals will be a subset of the progeny of heterozygous females, specifically those offspring homozygous for QG2082 alleles on the right arm of chromosome I or the center of chromosome III. Under a *peel* hypothesis, the only affected individuals will be the comparable subset of the progeny of heterozygous males. A detailed accounting of the expected results under the *Medea* hypothesis are shown in Figure S4.

Worms were maintained on OP50 *E. coli* at 25°C during the experiments and for three generations beforehand. For each cross, 9 L4 females and 10 L4 males were transferred to a 6cm plate on day 1 and left to mature and mate overnight. The following day (day 2), females with mating plugs were transferred to individual plates for a 7-hour egg lay period. Crosses were handled in a random order and plates were blinded to genotype for subsequent scoring. After the egg lay, females were then removed and the embryos laid on the plate were counted. On day 3, 18 hours after the end of the egg lay, unhatched embryos and deformed, arrested L1 larvae were counted. The deformed larvae were characteristically arrested in a two- or three-fold posture as if unable to move after hatching. On day 4, 42-48 hours after the end of the egg lay period, wild-type males and females, L4 males and females, and active L1-L3 larvae were counted. At this timepoint all larvae are the product of the original egg lay, as eggs laid by the adult offspring have not yet hatched. Raw count data are reported in Table S3.

To summarize the results, counts were pooled by cross and plotted as proportions, with the total being the number of embryos counted on day 2, immediately after the egg lay. In many cases, the sum of unhatched embryos, deformed L1s, developmentally delayed larvae, and wild-type adults was less than the number of embryos present initially. This gap is due in part to the difficulty in observing the small arrested L1 larvae, but it also reflects slight errors in counting each of the classes of embryos and worms. This gap between total embryos counted on day 2 and total offspring accounted for on days 3 and 4 is recorded as “missing” in Figure 3.

Under the two-*Medea* model, a greater proportion of QG2082-backcross progeny will be affected than F_2_ progeny. More specifically, assuming the *Medeas* have independent (*i.e,* multiplicative) effects, the difference in affected proportions between these two classes of progeny is an estimator of 1/16 *β*_I_ + 1/16 *β*_III_ + 3/16 *β*_I_*β*_III_, where the *β*s are the penetrances of the *Medea* loci on chromosomes I and III. Pooling observations across relevant experiments, we count 517/827 affected offspring in the QG2082 backcrosses and 644/1544 in the F_2_s. The difference in affected proportions is 0.21±0.04 (95% confidence interval calculated via Agresti-Caffo method). If the two *Medeas* have the same penetrance, the data provide an estimate of the penetrance as 0.77±0.10. Alternatively, if one *Medea* is completely penetrant, the data imply the other has a penetrance ∼0.59±0.16.

### Repetitive Element Analysis

We generated a redundant set of repetitive element libraries, identified *de novo* using the tools RepeatModeler (Flynn *et al.,* 2020), TransposonPSI (B. Haas, 2007), LTRharvest (Ellinghaus *et al.,* 2008), LTRdigest (Steinbiss *et al.,* 2009), SINE-Scan (Mao & Wang, 2017), SineFinder (Wenke *et al.,* 2011), TirVish (Gremme *et al.,* 2013), HelitronScanner (Xiong *et al.,* 2014), MITE-Tracker (Crescente *et al.,* 2018), MUSTv2 (Ge *et al.,* 2017), and MiteFinderII (Hu *et al.,* 2018). These libraries were merged with nematode-specific repeat libraries from Repbase (Bao *et al.,* 2015) and Dfam (Storer *et al.,* 2021) to create a single redundant repeat library. To remove redundancies, we used USEARCH (Edgar, 2010) to cluster the sequences in the library. The resulting non-redundant library was further filtered for any potential non-repeat clusters by using BlastX (Camacho et al., 2009) to search against nematode-specific proteins and non-coding RNAs.

The repeat discovery pipeline was adapted from a protocol previously used to characterize repeats in the *C. inopinata* genome (Coghlan *et al.,* 2018; Woodruff & Teterina, 2020). To validate this pipeline, we replicated previously reported repeat contents for *C. inopinata*, *C. elegans*, *C. briggsae*, *C. nigoni*, *C. remanei*, and *C. bovis* (Woodruff & Teterina, 2020). We used the hierarchical classification schema applied in other *Caenorhabditis* species (Woodruff & Teterina, 2020). The non-redundant repetitive element library was classified using multiple tools: RepeatClassifier (Flynn *et al.,* 2020), Dfam Classifier (Storer *et al.,* 2021), TransposonUltimate RFSB Classifier (Riehl *et al.,* 2021), and Geneious Sequence Classifier (https://www.geneious.com). We assigned the consensus classification from these tools, with conflicting classifications labeled as “unknown.”

Repeat libraries were generated independently for each genome assembly, and each genome was masked with RepeatMasker (http://www.repeatmasker.org/), using only their specific repeat libraries. We used the final RepeatMasker-generated masking results to characterize the genomic repeat landscape at both the global and taxonomic levels. To account for nested repeats, we disjoined the GFF so that each base pair in the genome was assigned to only one TE copy. This allowed us to distinguish between the number of repeat insertions and the number of bases covered by a given repeat taxon (Anderson *et al.,* 2019). We estimated the age of repetitive elements by calculating Kimura distances (Kapusta et al., 2017), which measure the divergence of individual repeat copies from their consensus sequence. Kimura distances were extracted from the “.align” file generated by RepeatMasker, and we calculated the mean Kimura distances across non-overlapping 10-kb windows for all repeat taxonomic ranks.

The repeat library, repeat classification, repeat gff, and tables of summary statistics contributing to figures S2 and S5, are provided as Supplementary File 4.

### Gene Structure

QG2082 worms were raised under standard conditions with OP50 *E. coli* food and then split into five conditions, as described in Sloat *et al*. (2022): 1) CemBio bacterial strains (Dirksen *et al.,* 2020) as food; 2) OP50 and standard conditions; 3) OP50 plus heat stress; 4) OP50 plus cold stress; 5) starvation. Temperature stresses consisted of 35°C or 4°C for 2 hours followed by a 2-hour recovery prior to RNA extraction. Total RNA was isolated using TRIzol following the protocol described in Green & Sambrook (2020). The mRNA libraries were constructed using the Illumina Stranded mRNA Prep Ligation protocol. These barcoded libraries were then pooled and sequenced using a NextSeq 500 MidOutput 2X150 for 300 cycles.

For the Iso-seq RNA extraction, we grew two sets of mixed-stage worms. One was well fed and the other starved for 2 days to generate a mix of dauer, arrested L1s, and starved adults. Worms were washed 5x in M9 to remove bacteria. The resulting pellet was resuspended in a mortar and pestle tube with 100 µL of TriZol and flash frozen in liquid nitrogen. The frozen powder was ground and 900 µL of additional TRIzol was added. The frozen lysate was then repeatedly (x10) thawed at 37°C on a heat block, and vortexed for 30 seconds. Then, 200 µl of chloroform was added per 1 ml TRIzol. The sample was mixed by inversion for 15 seconds before incubation at room temperature for 3 minutes for phase separation. The sample was then spun at 12,000 g for 15 mins at 4°C. The upper aqueous phase was then transferred to a fresh tube. We next added an equal volume of 100% ethanol, mixed by pipetting, and transferred the whole to a Qiagen RNeasy spin column, which was processed according to the manufacturer’s instructions. The resulting RNA was measured with the Qubit and the samples were combined to equal concentrations. The RNA was sent to the Center for Genomic and Computational Biology at Duke University for PacBio Sequel Sequencing.

We generated *de novo* gene models generally following the protocol of Doyle *et al*. (2020). RNAseq reads were mapped to the genome using STAR (Dobin & Gingeras, 2015). The resulting BAM was used as hints for BRAKER2 (Brůna *et al.,* 2021), which trains AUGUSTUS (Stanke *et al.,* 2006) using *ab initio* predictions generated by GeneMark-ES (Ter-Hovhannisyan *et al.,* 2008). Attempts to use mapped Iso-Seq or proteins of closely related Nematodes as input for BRAKER2 yielded poor results. Therefore, only short read RNAseq was used.

To better utilize the Iso-Seq data we mapped it to the genome using PASA (Haas *et al.,* 2003). We then loaded the BRAKER2 annotation into PASA to iteratively update existing annotations based on alignment evidence. We performed two rounds of updates to merge the two annotations following the recommendations of the PASA authors. We incorporated further sources of evidence by mapping *C. elegans* proteins to the *C. becei* genome using Exonerate (Slater & Birney, 2005) with an alignment score threshold of 50%. GFFs were subsequently extracted from the exonerate output with Exonerate_to_evm_gff3.pl from EvidenceModeler (Haas *et al.,* 2008). BRAKER2, PASA, and exonerate annotations were combined using weights of 2, 5, and 2 respectively in EvidenceModeler. The evidence modeler annotations were updated one last time for two rounds in PASA to incorporate additional evidence from long-read transcripts adding any UTR and isoform information that may have been missed or discarded.

### Genomic GC Content

We calculated the GC content with bedtools *nuc* (Quinlan & Hall, 2010). For analyses by functional class, we used species-specific GFF annotation files and computed base composition for each genomic feature. A nested masking approach was employed, where genomes were first masked for exons and subsequently for repeats, allowing for the separation of intergenic and intronic regions from exonic and repetitive sequences.

### Codon and Amino Acid Usage

To identify differences in codon preference among *Caenorhabditis* species, we analyzed the mean Relative Synonymous Codon Usage (RSCU) values (Sharp & Li, 1986). RSCU is a measure of codon bias, indicating how frequently a given codon is used relative to the expected frequency if all synonymous codons for an amino acid were used equally. An RSCU value of 1 suggests that a codon is used at its expected frequency under no bias, whereas values greater than 1 indicate codon preference, and values less than 1 indicate underrepresentation of that codon. Coding sequences for each species were retrieved from FASTA files, and RSCU values were calculated using the *uco* function from the *seqinr* package in R (Charif & Lobry, 2007).

Amino acid composition was calculated by mapping codons to their corresponding amino acids and summing codon counts for each amino acid across all coding sequences (CDS) in a species. Data represented in Figure 5 are provided in Supplementary File 5.

### Synteny Analysis

To visualize conserved gene order, we identified single-copy orthologs shared among *C. becei*, *C. niphades*, and *C. elegans* and then generated plots connecting the chromosomal positions of these orthologs across species. We found that chromosomes of *C. niphades* (Sun *et al.,* 2022) were syntenic with their corresponding chromosomes in *C. elegans* and *C. becei*, but the order of the genes along five of the chromosomes (I, II, III, V, and X) was largely inverted, suggesting that the genome fasta for these chromosomes represents the reverse complement relative to the other species. Chromosome IV, in contrast, was highly collinear between *C. becei* and *C. niphades*. For the plots in Figure 6, we reversed the orientation of the five *C. niphades* chromosomes to maximize the collinearity.

### Ortholog Identification & Functional Annotation

Predicted protein sequences for *C. becei* were generated from the reference genome and gene annotation using gffread v0.12.7 (Pertea & Pertea, 2020). Orthologous gene families were identified using OrthoFinder2 v2.5.4 (Emms & Kelly, 2019) from longest-isoform proteomes of *C. becei, C. briggsae, C. elegans, C. inopinata, C. latens, C. nigoni, C. niphades, C. remanei, C. sinica,* and *C. tropicalis*. Pairwise protein sequence similarity searches were performed with DIAMOND v2.0.15 (Buchfink et al., 2021) in ultra-sensitive mode, and orthogroups were inferred using the default OrthoFinder clustering procedure.

*C. becei* genes were classified according to their representation within orthogroups constructed from the ten *Caenorhabditis* proteomes. Genes were classified as duplicated when an orthogroup contained more than one *C. becei* gene. Genes were classified as single-copy orthologs when exactly one *C. becei* gene was assigned to an orthogroup, regardless of the number of copies present in the other species represented in that orthogroup. Genes that OrthoFinder did not assign to any orthogroup were classified as unassigned. Thus, the unassigned category reflects the absence of detectable orthogroup membership in this comparative analysis rather than definitive absence of homologs or orthologs in the other species.

Functional annotation was performed on the 22,965 retained longest-isoform *C. becei* protein sequences using InterProScan v5.65 (Blum et al., 2021), with Gene Ontology and pathway annotation enabled. Because InterProScan integrates predictions from multiple member databases, the output can contain several annotations describing the same or closely corresponding region of a protein. Annotation records were therefore filtered to reduce redundant representation of the same protein feature while retaining complementary annotations. Pfam, PANTHER, MobiDBLite, and Coils were treated as the primary annotation sources. Calls from other InterProScan analyses were retained as fallback annotations only for proteins lacking annotations from these primary sources.

Redundancy among overlapping primary-source annotations was further reduced using protein coordinates. When a Pfam annotation and a non-Pfam annotation on the same protein overlapped and both their start and end coordinates differed by no more than 10 amino acids, the Pfam annotation was retained and the corresponding non-Pfam annotation was removed. Non-Pfam annotations that did not meet this near-coincident overlap criterion were retained as potentially distinct protein features. Exact duplicate annotation records were also removed.

To facilitate biologically interpretable comparisons among genes, retained InterProScan annotations were grouped into broader curated domain-family or protein-feature categories. This reduced overly granular or database-specific labels while preserving distinctions among biologically different features. The curated classification used for downstream analyses is recorded in the domain_corrected field of File S2, while the underlying annotation information was retained, including the member-database source, signature accession and description, protein coordinates, InterPro accession and description when available, and associated GO annotations. Genes without a retained domain-family or protein-feature annotation were also retained in the final annotation dataset, ensuring that the complete set of 22,965 longest-isoform C. becei genes was represented, including genes for which no usable annotation was identified.

For gene-level domain-family analyses, each gene was counted at most once within a given domain family, even when that domain occurred multiple times within the same protein. Genes containing more than one distinct domain family were counted once in each relevant family. Domain-family counts therefore represent the number of genes containing each domain family, but the same gene can contribute to more than one domain-family count.

## ACKNOWLEDGMENTS

This work was supported by grants GM121828 and GM141906 from the National Institute for General Medical Sciences and HG013015 from National Human Genome Research Institute, and by support from the Zegar Foundation to the NYU GenCore facility. We thank the staff of NYU GenCore, the NYU IT High Performance Computing Team, Maggie Weitzman and the University of Oregon GC3 for HiFi data, Dovetail Genomics for Hi-C data, Duke University Genome Sequencing & Analysis Core for IsoSeq data, Arielle Martel for help in the lab, and the *Caenorhabditis* Genomes Project and WormBase for data. We thank Lewis Stevens and Asher Cutter for advice and suggestions.

## DATA AVAILABILITY STATEMENT

*C. becei* strain QG2082 genome assembly, and the HiFi, HiC, Illumina RNAseq, and PacBio Isoseq data used to assemble and annotate it, are associated with BioProject PRJNA989223.

## SUPPLEMENTARY FILES

**File S1. Genome annotation.** This zip-compressed gff3 file contains the genome annotation, including the 24,025 transcripts representing 22,965 unique genes, along with 122,867 repeat regions.

**File S2. Protein domain annotations.** This zip-compressed tsv file contains domain and protein-feature annotations for the complete set of 22,965 retained longest-isoform *C. becei* genes used in the orthology analyses. The table contains 82,376 rows because genes can have multiple retained annotation records, including different domain families or multiple occurrences of the same domain within a protein. Genes without a retained domain annotation are also included so that the table represents the complete analyzed gene set rather than only genes with InterProScan annotations.

The table column headings (shown `fixed width`) are as follows. `GENE_ID` gives the *C. becei* gene identifier, and `MRNA_ID` gives the identifier of the retained (longest isoform) transcript corresponding to that gene. `OG_id` gives the orthogroup identifier associated with the gene, and `OG_class` gives its orthology classification as single_copy_ortholog, duplicated_gene, or unassigned.

`analysis` identifies the InterProScan analysis or member database that produced the annotation record. Analyses represented in the table include Pfam, PANTHER, MobiDBLite, Coils, Gene3D, SUPERFAMILY, ProSiteProfiles, SMART, ProSitePatterns, CDD, PRINTS, and FunFam. A value of “-” is used for added records representing genes without a retained annotation. `signature_accession` gives the accession or identifier of the specific signature producing the feature call, and `signature_description` gives the corresponding signature description when available. `domain_corrected` gives the standardized domain-family or protein-feature label used in downstream domain analyses. This column consolidates annotations into the labels used for gene-level domain-family summaries; a value of “-” indicates that no usable domain label was assigned to that record.

`gene_chr`, `gene_start`, `gene_end`, and `gene_strand` give the chromosome, genomic start coordinate, genomic end coordinate, and strand of the gene, respectively. `mrna_chr`, `mrna_start`, `mrna_end`, and `mrna_strand` provide the corresponding chromosome, genomic coordinates, and strand for the retained representative transcript.

`domain_start` and `domain_end` give the start and end coordinates of the detected domain or protein feature within the translated protein sequence, in amino-acid coordinates. These values are absent for added records representing genes without a retained domain annotation. `interpro_accession` gives the integrated InterPro accession associated with a signature when one is available, and `interpro_description` gives the corresponding InterPro description. A value of “-” indicates that no corresponding InterPro entry was assigned. `go_annotations` contains Gene Ontology terms associated with the annotation record when provided by InterProScan; records without associated GO terms contain “-”.

**File S3. Reporter plasmid sequences.** Fasta-formatted DNA sequences for pSAS02 and pSAS06, plasmids for expression of male-specific GFP and female specific mCherry.

**File S4. Repeat annotation data.** This zip-compressed directory contains four files:

1. becei_Repeat_Taxonomy.xlsx describes the repeat classification scheme used in this study. `Fasta_header_classification` gives the classification used in the repeat-library FASTA headers, `RM_classification` gives the corresponding RepeatMasker classification, and `Class`, `Subclass`, `Order`, `Superfamily`, and `Family` give the hierarchical taxonomic assignments for each repeat type. Some repeat types do not have assignments at every taxonomic level.
2. becei_repeat_library.fa contains the nonredundant *C. becei* repeat library of 1,043 repeat-cluster sequences. FASTA headers contain the repeat-cluster identifier followed by its repeat classification.
3. becei_global_repeat_density_10kb_win_norm_dist_cent.tsv reports total repeat coverage in 10 kb windows across chromosomes I-V and X. `Chr` gives the chromosome, `BP` gives the starting coordinate of the window, `num_rep` gives the number of base pairs within the window annotated as repetitive, `species` gives the species identifier, and `norm_dist_center` gives the normalized distance of the window from the center of its chromosome, with values near 0 at the chromosome center and approaching 0.5 toward chromosome ends.
4. all_models_superfamily.tsv reports repeat abundance and chromosome-position analyses for individual repeat categories. `rep_class` identifies the repeat category and `rep_count` gives the number of repeat annotations assigned to that category. `adj_r2`, `F`, `numdf`, `dendf`, `beta_Estimate`, `beta_std_error`, `t_value`, and `p_value` report statistics from linear models testing the relationship between repeat abundance and normalized distance from the chromosome center. `arm_lower`, `arm_mean`, and `arm_upper` give the lower 95% confidence limit, mean, and upper 95% confidence limit for repeat abundance in chromosome arms, respectively, while `center_lower`, `center_mean`, and `center_upper` give the corresponding values for chromosome centers. `arm_cen_effect_size_lower`, `arm_cen_effect_size`, and `arm_cen_effect_size_upper` give the lower 95% confidence limit, Cohen’s *d* effect size, and upper 95% confidence limit for the comparison between chromosome arms and centers. `percent_genome_repeat` gives the percentage of the genome occupied by each repeat category, and `repeat_total_bp` gives the corresponding number of genomic base pairs. `p_adjust` gives the Benjamini-Hochberg-adjusted P value and `p_sig` indicates whether the association remains significant after multiple-testing correction. `species` gives the species identifier. `sort`, `label`, and `for_plot` are variables used for ordering and labeling repeat categories in Figure S2.

**File S5. Codon and amino acid usage data.** This .xlsx file contains three tabs, containing the numerical data plotted in Figure 5.

## SUPPLEMENTARY TABLES

**Table S1:** The genetic map of *C. becei*, with physical positions of markers on the genome assembly.

**Table S2.** Recombination domains in *C. becei* and other species

From genetic and physical maps for each chromosome, after excluding the terminal megabase from each end, we used segmented linear regression to identify three recombination-rate domains. The table records the positions of the boundaries (LC and CR, boundaries between the left arm and center and between the center and right arm, respectively), the percent of chromosome length that is in the left arm and tip, center domain, and right arm and tip, and then the recombination rates estimated for each domain from the regression slopes. Note that four of the 36 chromosome maps (6 chromosomes x 6 species) have recombination rate patterns that do not match the expectations of the regression, and the numbers in the table for these are not meaningful. These chromosomes – *C. tropicalis* X, *C. remanei* IV and X, and *P. pacificus* X – are indicated by “No” in the “Domains” column in the table. The *P. pacificus* X is likely affected by segregating inversions and so may not reflect the meiotic map in structural homozygotes.

**Table S3.** *Medea* data. Results of experimental crosses testing for Medea or Peel activity. These are the data underlying Figure 3B. Each row represents one worm’s progeny, indexed in column 1 to its randomly assigned plate number. Columns 2-10 count the numbers of embryos observed after egg lay, unhatched embryos and deformed larvae the next day, and then wild-type adult females, wild-type adult males, L4 females, L4 males, and L1-3 larvae on the subsequent day. Columns 11 and 12 record the genotypes of the parents of the cross, and column 13 records an identifier for which of the 16 classes of cross the row represents.

## REFERENCES

Adams, P. E., Crist, A. B., Young, E. M., Willis, J. H., Phillips, P. C., & Fierst, J. L. (2022). Slow recovery from inbreeding depression generated by the complex genetic architecture of segregating deleterious mutations. Molecular Biology and Evolution, 39, msab330

Aharonoff, A., Kim, J., Washington, A., & Ercan, S. (2025) Parallel Evolution of X Chromosome-Specific Structural Maintenance of Chromosomes Complexes in Two Nematode Lineages, Molecular Biology and Evolution 42 msaf270.

Al Arab, M., Höner Zu Siederdissen, C., Tout, K., Sahyoun, A.H., Stadler, P.F., & Bernt, M. (2017).Accurate annotation of protein-coding genes in mitochondrial genomes. Mol Phylogenet Evol. 106:209–216.

Andersen, E. C., & Rockman, M. V. (2022). Natural genetic variation as a tool for discovery in *Caenorhabditis* nematodes. Genetics 220, iyab156

Anderson, S. N., Stitzer, M. C., Brohammer, A. B., Zhou, P., Noshay, J. M., O’Connor, C. H., Hirsch, C. D., Ross-Ibarra, J., Hirsch, C. N., & Springer, N. M. (2019). Transposable elements contribute to dynamic genome content in maize. The Plant Journal 100, 1052–1065.

Andolfatto, P., Davison, D., Erezyilmaz, D., Hu, T. T., Mast, J., Sunayama-Morita, T., & Stern, D. L. (2011). Multiplexed shotgun genotyping for rapid and efficient genetic mapping. Genome Research, 21, 610–617.

Baer, C. F., Joyner-Matos, J., Ostrow, D., Grigaltchik, V., Salomon, M. P., & Upadhyay, A. (2010). Rapid decline in fitness of mutation accumulation lines of gonochoristic (outcrossing) *Caenorhabditis* nematodes. Evolution 64, 3242–3253.

Bao, W., Kojima, K. K., & Kohany, O. (2015). Repbase Update, a database of repetitive elements in eukaryotic genomes. Mobile DNA 6, 11.

Barnes, T. M., Kohara, Y., Coulson, A., & Hekimi, S. (1995). Meiotic recombination, noncoding DNA and genomic organization in *Caenorhabditis elegans*. Genetics, 141(1), 159–179.

Barrière, A., & Félix, M.-A. (2005). High local genetic diversity and low outcrossing rate in *Caenorhabditis elegans* natural populations. Current Biology 15, 1176–1184.

Beeman, R. W., Friesen, K. S., & Denell, R. E. (1992). Maternal-effect selfish genes in flour beetles. Science 256, 89–92.

Ben-David, E., Burga, A., & Kruglyak, L. (2017). A maternal-effect selfish genetic element in *Caenorhabditis elegans*. Science 356, 1051–1055.

Ben-David, E., Pliota, P., Widen, S. A., Koreshova, A., Lemus-Vergara, T., Verpukhovskiy, P., Mandali, S., Braendle, C., Burga, A., & Kruglyak, L. (2021). Ubiquitous selfish toxin-antidote elements in *Caenorhabditis* species. Current Biology 31, 990–1001.

Blum, M., Chang, H.-Y., Chuguransky, S., Grego, T., Kandasaamy, S., Mitchell, A., Nuka, G., Paysan-Lafosse, T., Qureshi, M., Raj, S., Richardson, L., Salazar, G. A., Williams, L., Bork, P., Bridge, A., Gough, J., Haft, D. H., Letunic, I., Marchler-Bauer, A., … Finn, R. D. (2021). The InterPro protein families and domains database: 20 years on. Nucleic Acids Research, 49, D344–D354.

Brůna, T., Hoff, K. J., Lomsadze, A., Stanke, M., & Borodovsky, M. (2021). BRAKER2: automatic eukaryotic genome annotation with GeneMark-EP+ and AUGUSTUS supported by a protein database. NAR Genomics and Bioinformatics, 3, lqaa108.

Buchfink, B., K. Reuter, & H.-G. Drost. (2021). Sensitive protein alignments at tree-of-life scale using DIAMOND. Nature Methods 18, 366–368.

Camacho, C., Coulouris, G., Avagyan, V., Ma, N., Papadopoulos, J., Bealer, K., & Madden, T. L. (2009). BLAST+: architecture and applications. BMC Bioinformatics 10, 421.

Capella-Gutiérrez, S., Silla-Martínez, J. M., & Gabaldón, T. (2009). trimAl: a tool for automated alignment trimming in large-scale phylogenetic analyses. Bioinformatics 25, 1972–1973.

Carelli, F. N., Cerrato, C., Dong, Y., Appert, A., Dernburg, A., & Ahringer, J. (2022). Widespread transposon co-option in the *Caenorhabditis* germline regulatory network. Science Advances 8, eabo4082.

Charif, D., & Lobry, J. R. (2007). SeqinR 1.0-2: A contributed package to the R project for statistical computing devoted to biological sequences retrieval and analysis. In Structural Approaches to Sequence Evolution (pp. 207–232). Springer Berlin Heidelberg.

Cheng H, Concepcion GT, Feng X, Zhang H, Li H. (2021) Haplotype-resolved *de novo* assembly using phased assembly graphs with hifiasm. Nat Methods 18, 170–175.

Coghlan, A., Coghlan, A., Tsai, I. J., & Berriman, M. (2018). Creation of a comprehensive repeat library for a newly sequenced parasitic worm genome. Protocol Exchange. 10.1038/protex.2018.054

Crescente, J. M., Zavallo, D., Helguera, M., & Vanzetti, L. S. (2018). MITE Tracker: an accurate pproach to identify miniature inverted-repeat transposable elements in large genomes. BMC Bioinformatics, 19, 348.

Csankovszki, G., McDonel, P., & Meyer, B. J. (2004). Recruitment and spreading of the *C. elegans* dosage compensation complex along X chromosomes. Science 303, 1182–1185.

Cutter, A. D. (2018). X exceptionalism in *Caenorhabditis* speciation. Molecular Ecology 27, 3925–3934.

Cutter, A. D., & Charlesworth, B. (2006). Selection intensity on preferred codons correlates with overall codon usage bias in *Caenorhabditis remanei*. Current Biology 16, 2053–2057

Cutter, A.D., Wasmuth, J.D. & Blaxter, M.L. (2006). The evolution of biased codon and amino acid usage in nematode genomes. Molecular biology and evolution, 23(12), pp.2303–2315.

Cutter, A.D., Wasmuth, J.D. & Washington, N.L. (2008). Patterns of molecular evolution in *Caenorhabditis* preclude ancient origins of selfing. Genetics, 178(4), pp.2093–2104.

Danecek P, Bonfield JK, Liddle J, Marshall J, Ohan V, Pollard MO, Whitwham A, Keane T, McCarthy SA, Davies RM, Li H. (2021) Twelve years of SAMtools and BCFtools. Gigascience 10, giab008.

Daul, A. L., Andersen, E. C., & Rougvie, A. E. (2019). The *Caenorhabditis* Genetics Center (CGC) and the *Caenorhabditis elegans* Natural Diversity Resource. In The Biological Resources of Model Organisms (pp. 69–94). CRC Press.

Dayi, M., Kanzaki, N., Sun, S., Ide, T., Tanaka, R., Masuya, H., Okabe, K., Kajimura, H., & Kikuchi, T. (2021). Additional description and genome analyses of *Caenorhabditis auriculariae* representing the basal lineage of genus *Caenorhabditis*. Scientific Reports, 11 6720.

Dirksen, P., Assié, A., Zimmermann, J., Zhang, F., Tietje, A.-M., Marsh, S. A., Félix, M.-A., Shapira, M., Kaleta, C., Schulenburg, H., & Samuel, B. S. (2020). CeMbio - The *Caenorhabditis elegans* Microbiome Resource. G3 10, 3025–3039.

Dobin, A., & Gingeras, T. R. (2015). Mapping RNA-seq reads with STAR. Current Protocols in Bioinformatics 51, 11–14.

Dolgin, E. S., Charlesworth, B., Baird, S. E., & Cutter, A. D. (2007). Inbreeding and outbreeding depression in *Caenorhabditis* nematodes. Evolution 61, 1339–1352.

Donath, A., Jühling, F., Al-Arab, M., Bernhart, S.H., Reinhardt, F., Stadler, P.F., Middendorf, M., & Bernt, M. (2019). Improved annotation of protein-coding genes boundaries in metazoan mitochondrial genomes. Nucleic Acids Res. 47, 10543–10552.

Doyle, S. R., Tracey, A., Laing, R., Holroyd, N., Bartley, D., Bazant, W., Beasley, H., Beech, R., Britton, C., Brooks, K., Chaudhry, U., Maitland, K., Martinelli, A., Noonan, J. D., Paulini, M., Quail, M. A., Redman, E., Rodgers, F. H., Sallé, G., … Cotton, J. A. (2020). Genomic and transcriptomic variation defines the chromosome-scale assembly of *Haemonchus contortus*, a model gastrointestinal worm. Communications Biology 3, 656.

Durand NC, Shamim MS, Machol I, Rao SS, Huntley MH, Lander ES, & Aiden EL. (2016). Juicer provides a one-click system for analyzing loop-resolution Hi-C experiments. Cell Syst. 3, 95–8.

Edgar, R. C. (2010). Search and clustering orders of magnitude faster than BLAST. Bioinformatics 26, 2460–2461.

Ellinghaus, D., Kurtz, S., & Willhoeft, U. (2008). LTRharvest, an efficient and flexible software for *de novo* detection of LTR retrotransposons. BMC Bioinformatics 9, 18.

Emms, D. M., & Kelly, S. (2019). OrthoFinder: phylogenetic orthology inference for comparative genomics. Genome Biology 20, 238.

Eurmsirilerd, E., & Maduro, M. F. (2020). Evolution of developmental GATA factors in nematodes. Journal of Developmental Biology 8, 27.

Farboud, B., Nix, P., Jow, M. M., Gladden, J. M., & Meyer, B. J. (2013). Molecular antagonism between X-chromosome and autosome signals determine nematode sex. Genes and Development 27, 1159–1178.

Félix, M.-A., Braendle, C., & Cutter, A. D. (2014). A streamlined system for species diagnosis in *Caenorhabditis* (Nematoda: Rhabditidae) with name designations for 15 distinct biological species. PloS One 9, e94723.

Félix, M.-A., Jovelin, R., Ferrari, C., Han, S., Cho, Y. R., Andersen, E. C., Cutter, A. D., & Braendle, C. (2013). Species richness, distribution and genetic diversity of *Caenorhabditis* nematodes in a remote tropical rainforest. BMC Evolutionary Biology 13, 10.

Ferrari, C., Salle, R., Callemeyn-Torre, N., Jovelin, R., Cutter, A. D., & Braendle, C. (2017). Ephemeral-habitat colonization and neotropical species richness of *Caenorhabditis* nematodes. BMC Ecology 17, 43.

Flynn, J. M., Hubley, R., Goubert, C., Rosen, J., Clark, A. G., Feschotte, C., & Smit, A. F. (2020). RepeatModeler2 for automated genomic discovery of transposable element families. Proc Natl Acad Sci USA 117, 9451–9457.

Foster, P.G., Jermiin, L.S. and Hickey, D.A., 1997. Nucleotide composition bias affects amino acid content in proteins coded by animal mitochondria. Journal of Molecular Evolution 44, 282–288.

Fusca, D. D., Kasimatis, K.R., Zhu, H. V., & Cutter, A.D. 2025. Dynamic birth and death of Argonaute gene family functional repertoire across *Caenorhabditis* nematodes. Genome Biology and Evolution 17, evaf106.

Fusca, D. D., Dall’Acqua, M. N., Sánchez-Ramirez, S., & Cutter, A. D. 2025. Phylogenomic timetree-calibrated speciation clocks for *Caenorhabditis* nematodes reveal slow but disproportionate accumulation of post-zygotic reproductive isolation. PLoS Genetics 21, e1011852.

Ge, R., Mai, G., Zhang, R., Wu, X., Wu, Q., & Zhou, F. (2017). MUSTv2: an improved *de novo* detection program for recently active Miniature Inverted Repeat Transposable Elements (MITEs). Journal of Integrative Bioinformatics 14, 20170029.

Gladden, J. M., & Meyer, B. J. (2007). A ONECUT Homeodomain protein communicates X chromosome dose to specify *Caenorhabditis elegans* sexual fate by repressing a sex switch gene. Genetics 177, 1621–1637.

Gonzalez de la Rosa, P. M., Thomson, M., Trivedi, U., Tracey, A., Tandonnet, S., & Blaxter, M. (2021). A telomere-to-telomere assembly of *Oscheius tipulae* and the evolution of rhabditid nematode chromosomes. G3 11, 1–17.

Green, M. R., & Sambrook, J. (2020). Total RNA extraction from *Caenorhabditis elegans*. Cold Spring Harbor Protocols 2020, 101683.

Gremme, G., Steinbiss, S., & Kurtz, S. (2013). GenomeTools: a comprehensive software library for efficient processing of structured genome annotations. IEEE/ACM Transactions on Computational Biology and Bioinformatics 10, 645–656.

Haas, B. (2007). TransposonPSI: an application of PSI-Blast to mine (retro-) transposon ORF homologies. Broad Institute, Cambridge, MA, USA. https://transposonpsi.sourceforge.net/

Haas, B. J., Delcher, A. L., Mount, S. M., Wortman, J. R., Smith, R. K., Jr, Hannick, L. I., Maiti, R., Ronning, C. M., Rusch, D. B., Town, C. D., Salzberg, S. L., & White, O. (2003). Improving the *Arabidopsis* genome annotation using maximal transcript alignment assemblies. Nucleic Acids Research 31, 5654–5666.

Haas, B. J., Salzberg, S. L., Zhu, W., Pertea, M., Allen, J. E., Orvis, J., White, O., Buell, C. R., & Wortman, J. R. (2008). Automated eukaryotic gene structure annotation using EVidenceModeler and the Program to Assemble Spliced Alignments. Genome Biology 9, R7.

Harbin, J.P., Shen, Y., Abubakar, A.H. and Ellis, R.E., 2026. Haldane’s law works through X: Autosome incompatibility in *Caenorhabditis briggsae/C. nigoni* hybrids. Nature communications 17, 1679.

Huang, Y., Lo, Y.-H., Hsu, J.-C., Le, T. S., Yang, F.-J., Chang, T., Braendle, C., & Wang, J. (2023). Widespread sex ratio polymorphism in *Caenorhabditis* nematodes. Royal Society Open Science 10, 221636.

Hu, J., Zheng, Y., & Shang, X. (2018). MiteFinderII: a novel tool to identify miniature inverted-repeat transposable elements hidden in eukaryotic genomes. BMC Medical Genomics 11(Suppl 5), 101.

Jhaveri, N., Gupta, B. and Chamberlin, H.M., 2025. Coincident evolution and functional adaptation of the taxonomically restricted genes *ivph-3* and *gon-14* in *Caenorhabditis* nematodes. Biology Open 14, bio062018.

Johnson, A. J., & Gomez, D. F. (2020). General assembly and alignment in Geneious v1. 10.17504/protocols.io.bnvfme3n

Kapusta, A., Suh, A., & Feschotte, C. (2017). Dynamics of genome size evolution in birds and mammals. Proc Natl Acad Sci USA 114, E1460–E1469.

Katoh, K., & Standley, D. M. (2013). MAFFT multiple sequence alignment software version 7: improvements in performance and usability. Molecular Biology and Evolution 30, 772–780.

Kiontke, K. C., Félix, M.-A., Ailion, M., Rockman, M. V., Braendle, C., Pénigault, J.-B., & Fitch, D. H. A. (2011). A phylogeny and molecular barcodes for *Caenorhabditis*, with numerous new species from rotting fruits. BMC Evolutionary Biology 11, 339.

Kiontke, K., Gavin, N. P., Raynes, Y., Roehrig, C., Piano, F., & Fitch, D. H. A. (2004). *Caenorhabditis* phylogeny predicts convergence of hermaphroditism and extensive intron loss. Proc Natl Acad Sci USA 101, 9003–9008.

Knight, R.D., Freeland, S.J. and Landweber, L.F., 2001. A simple model based on mutation and selection explains trends in codon and amino-acid usage and GC composition within and across genomes. Genome biology 2, research0010-1.

Kolmogorov, M., Yuan, J., Lin, Y., & Pevzner, P.A. (2019). Assembly of long, error-prone reads using repeat graphs. Nat Biotechnol. 37, 540–546.

Kursel, L. E., Cope, H. D., & Rog, O. (2021). Unconventional conservation reveals structure-function relationships in the synaptonemal complex. bioRxiv. 10.1101/2021.06.16.448737

Lamelza, P., Young, J. M., Noble, L. M., Caro, L., Isakharov, A., Palanisamy, M., Rockman, M. V., Malik, H. S., & Ailion, M. (2019). Hybridization promotes asexual reproduction in *Caenorhabditis* nematodes. PLoS Genetics 15, e1008520.

Large, C.R., Khanal, R., Hillier, L., Huynh, C., Kubo, C., Kim, J., Waterston, R.H. and Murray, J.I., 2025. Lineage-resolved analysis of embryonic gene expression evolution in *C. elegans* and *C. briggsae*. Science 388, eadu8249.

Le, S. Q., & Gascuel, O. (2008). An improved general amino acid replacement matrix. Molecular Biology and Evolution 25, 1307–1320.

Le, T. S., Yang, F.-J., Lo, Y.-H., Chang, T. C., Hsu, J.-C., Kao, C.-Y., & Wang, J. (2017). Non-Mendelian assortment of homologous autosomes of different sizes in males is the ancestral state in the *Caenorhabditis* lineage. Scientific Reports 7, 12819.

Lee, D., Zdraljevic, S., Stevens, L., Wang, Y., Tanny, R. E., Crombie, T. A., Cook, D. E., Webster, A. K., Chirakar, R., Baugh, L. R., Sterken, M. G., Braendle, C., Felix, M.-A., Rockman, M. V., and Andersen, E. C. (2021). Balancing selection maintains hyper-divergent haplotypes in *Caenorhabditis elegans*. Nature Eco. Evo. 5, 795–807.

Levya-Díaz, E., & Hobert, O. (2022). Robust regulatory architecture of pan-neuronal gene expression. Current Biology 32, 1715–1727.

Li, H. (2018) Minimap2: pairwise alignment for nucleotide sequences, Bioinformatics 34, 3094–3100.

Li, Y., Gao, Y., Ma, J., Gao, Y., Tang, J., Yang, R., Wang, J., Zhou, W., Zhang, H., Shao, W. & Liu, Z. (2025). Aberrant X chromosome dosage compensation causes hybrid male inviability in *Caenorhabditis*. Proceedings of the National Academy of Sciences 122, e2507166122.

Libuda, D. E., Uzawa, S., Meyer, B. J., & Villeneuve, A. M. (2013). Meiotic chromosome structures constrain and respond to designation of crossover sites. Nature 502, 703–706.

Maciejowski, J., Ahn, J. H., Cipriani, P. G., Killian, D. J., Chaudhary, A. L., Lee, J. I., Voutev, R., Johnsen, R. C., Baillie, D. L., Gunsalus, K. C., Fitch, D. H. A., & Hubbard, E. J. A. (2005). Autosomal genes of autosomal/X-linked duplicated gene pairs and germ-line proliferation in *Caenorhabditis elegans*. Genetics 169, 1997–2011.

Maduro, M. F. (2020). Evolutionary dynamics of the SKN-1 → MED → END-1,3 regulatory gene cascade in *Caenorhabditis* endoderm specification. G3 10, 333–356.

Mao, H., & Wang, H. (2017). SINE_scan: an efficient tool to discover short interspersed nuclear elements (SINEs) in large-scale genomic datasets. Bioinformatics 33, 743–745.

Memar, N., Schiemann, S., Hennig, C., Findeis, D., Conradt, B., & Schnabel, R. (2019). Twenty million years of evolution: The embryogenesis of four *Caenorhabditis* species are indistinguishable despite extensive genome divergence. Developmental Biology 447, 182–199.

Meyer, B. (2022). Mechanisms of sex determination and X-chromosome dosage compensation. Genetics 220, iyab197.

Moya, N.D., Yan, S.M., McCoy, R.C., & Andersen, E.C. (2025). The long and short of hyperdivergent regions. Trends in Genetics 41, 303–314.

Muggeo, V. M. R. (2008). Segmented: An R package to fit regression models with broken-line relationships. R News 8, 20–25.

Nelson, C., & Ambros, V. (2021). A cohort of *Caenorhabditis* species lacking the highly conserved let-7 microRNA. G3 11, jkab022

Nguyen, L.-T., Schmidt, H. A., von Haeseler, A., & Minh, B. Q. (2015). IQ-TREE: a fast and effective stochastic algorithm for estimating maximum-likelihood phylogenies. Molecular Biology and Evolution 32, 268–274.

Noble, L. M., Rockman, M. V., & Teotónio, H. (2021). Gene-level quantitative trait mapping in *Caenorhabditis elegans*. G3 11, jkaa061

Noble, L. M., Yuen, J., Stevens, L., Moya, N., Persaud, R., Moscatelli, M., Jackson, J. L., Zhang, G., Chitrakar, R., Baugh, L. R., Braendle, C., Andersen, E. C., Seidel, H. S., & Rockman, M. V. (2021). Selfing is the safest sex for *Caenorhabditis tropicalis*. eLife 10, e62587.

Nurk S, Walenz BP, Rhie A, Vollger MR, Logsdon GA, Grothe R, Miga KH, Eichler EE, Phillippy AM, Koren S. (2019). HiCanu: accurate assembly of segmental duplications, satellites, and allelic variants from high-fidelity long reads. Genome Res. 30, 1291–1305.

Paradis, E., & Schliep, K. (2019). ape 5.0: an environment for modern phylogenetics and evolutionary analyses in R. Bioinformatics 35, 526–528.

Parée, T., Noble, L., Roze, D., & Teotónio, H. (2025). Selection can favor a recombination landscape that limits polygenic adaptation. Molecular Biology and Evolution 42, msae273

Pertea, G., & M. Pertea. (2020). GFF Utilities: GffRead and GffCompare. F1000Res doi 10.12688/f1000research.23297.2.

Poplin, R., Chang, P.C., Alexander, D., Schwartz, S., Colthurst, T., Ku, A., Newburger, D., Dijamco, J., Nguyen, N., Afshar, P.T., Gross, S.S., Dorfman, L., McLean, C.Y., & DePristo, M.A. (2018). A universal SNP and small-indel variant caller using deep neural networks. Nat Biotechnol 36, 983–987.

Quinlan, A. R., & Hall, I. M. (2010). BEDTools: a flexible suite of utilities for comparing genomic features. Bioinformatics 26, 841–842.

R Core Team. (2024) R: A Language and Environment for Statistical Computing. R Foundation for Statistical Computing, Vienna, Austria.

Reilly, M. B., Cros, C., Varol, E., Yemini, E., & Hobert, O. (2020). Unique homeobox codes delineate all the neuron classes of *C. elegans*. Nature 584, 595–601.

Riehl, K., Riccio, C., Miska, E. A., & Hemberg, M. (2022). TransposonUltimate: software for transposon classification, annotation and detection. Nucleic Acids Research 50, gkac136.

Rillo-Bohn, R., Adilardi, R., Mitros, T., Avşaroğlu, B., Stevens, L., Köhler, S., Bayes, J., Wang, C., Lin, S., Baskevitch, K. A., Rokhsar, D. S., & Dernburg, A. F. (2021). Analysis of meiosis in *Pristionchus pacificus* reveals plasticity in homolog pairing and synapsis in the nematode lineage. eLife 10, e70990.

Rockman, M.V., Tintori, S.C., Nguyen, T.H. and Yomai, V.H. (2026). *Caenorhabditis* diversity on Pohnpei, Micronesia, provides evidence that the Elegans Supergroup has its roots in the Americas and diversified in the Pacific *en route* to Asia. Evolution 80, 1013–1034.

Rockman, M. V., Bernstein, M. B., Caglar, D., Cattani, M. V., Chang, A. S., Kaur, T, Noble, L. M., & Paaby, A. B. (2025). Variation in inbreeding depression within and among *Caenorhabditis* species. bioRxiv doi: 10.1101/2025.06.04.657873.

Rockman, M. V. (2025). Parental-effect gene-drive elements under partial selfing, or why do *Caenorhabditis* genomes have hyperdivergent regions? Genetics 229, iyae175.

Rockman, M. V., & Kruglyak, L. (2009). Recombinational landscape and population genomics of *Caenorhabditis elegans*. PLoS Genetics 5, e1000419.

Ross, J. A., Koboldt, D. C., Staisch, J. E., Chamberlin, H. M., Gupta, B. P., Miller, R. D., Baird, S. E., & Haag, E. S. (2011). *Caenorhabditis briggsae* recombinant inbred line genotypes reveal inter-strain incompatibility and the evolution of recombination. PLoS Genetics 7, e1002174.

Seidel, H. S., Rockman, M. V., & Kruglyak, L. (2008). Widespread genetic incompatibility in *C. elegans* maintained by balancing selection. Science 319, 589–594.

Seppey, M., Manni, M., & Zdobnov, E. M. (2019). BUSCO: Assessing genome assembly and annotation completeness. Methods in Molecular Biology 1962, 227–245.

Sharp, P. M., & Li, W. H. (1986). An evolutionary perspective on synonymous codon usage in unicellular organisms. Journal of Molecular Evolution 24, 28–38.

Shakya, P., Maulana, M. I., Danchin, E. G. J., Voogt, M. L., van de Ruitenbeek, S. J. S., Gimeno, J., Taranto, A. P., Blundell, A. C., Despot-Slade, E., Mestrovic, N., Mota, A. Z., Dai, D., Williamson, V. M., Sterken, M. G., & Siddique, S. (2025). High-resolution genome assembly and linkage mapping in *Meloidogyne hapla* reveal non-canonical telomere repeats and recombination hotspots associated with effector proteins. PLoS Pathogens 21, e1013706.

Shaw, C. L., & Kennedy, D. A. (2022). Developing an empirical model for spillover and emergence: Orsay virus host range in *Caenorhabditis*. Proceedings Biological Sciences 289, 20221165.

Slater, G. S. C., & Birney, E. (2005). Automated generation of heuristics for biological sequence comparison. BMC Bioinformatics 6, 31.

Sloat, S. A., Noble, L. M., Paaby, A. B., Bernstein, M., Chang, A., Kaur, T., Yuen, J., Tintori, S. C., Jackson, J. L., Martel, A., Salome Correa, J. A., Stevens, L., Kiontke, K., Blaxter, M., & Rockman, M. V. (2022). *Caenorhabditis* nematodes colonize ephemeral resource patches in neotropical forests. Ecology and Evolution 12, e9124.

Slos, D., Sudhaus, W., Stevens, L., Bert, W., & Blaxter, M. (2017). *Caenorhabditis monodelphis sp. n*.: defining the stem morphology and genomics of the genus *Caenorhabditis*. BMC Zoology 2, 4.

Snoek, B. L., Volkers, R. J. M., Nijveen, H., Petersen, C., Dirksen, P., Sterken, M. G., Nakad, R., Riksen, J. A. G., Rosenstiel, P., Stastna, J. J., Braeckman, B. P., Harvey, S. C., Schulenburg, H., & Kammenga, J. E. (2019). A multi-parent recombinant inbred line population of *C. elegans* allows identification of novel QTLs for complex life history traits. BMC Biology 17, 24.

Stanke, M., Keller, O., Gunduz, I., Hayes, A., Waack, S., & Morgenstern, B. (2006). AUGUSTUS: *ab initio* prediction of alternative transcripts. Nucleic Acids Research 34, W435–W439.

Steinbiss, S., Willhoeft, U., Gremme, G., & Kurtz, S. (2009). Fine-grained annotation and classification of *de novo* predicted LTR retrotransposons. Nucleic Acids Research 37, 7002–7013.

Stevens, L., Félix, M.-A., Beltran, T., Braendle, C., Caurcel, C., Fausett, S., Fitch, D., Frézal, L., Gosse, C., Kaur, T., Kiontke, K., Newton, M. D., Noble, L. M., Richaud, A., Rockman, M. V., Sudhaus, W., & Blaxter, M. (2019). Comparative genomics of 10 new *Caenorhabditis* species. Evolution Letters 3, 217–236.

Stevens, L., Moya, N. D., Tanny, R. E., Gibson, S. B., Tracey, A., Na, H., Chitrakar, R., Dekker, J., Walhout, A. J. M., Baugh, L. R., & Andersen, E. C. (2022). Chromosome-level reference genomes for two strains of *Caenorhabditis briggsae*: An improved platform for comparative genomics. Genome Biology and Evolution 14, evac042.

Stevens, L., Rooke, S., Falzon, L. C., Machuka, E. M., Momanyi, K., Murungi, M. K., Njoroge, S. M., Odinga, C. O., Ogendo, A., Ogola, J., Fèvre, E. M., & Blaxter, M. (2020). The genome of *Caenorhabditis bovis*. Current Biology 30, 1023–1031.

Stevens, L., Martinez-Ugalde, I., King, E., Wagah, M., Absolon, D., Bancroft, R., Gonzalez de la Rosa, P., Hall, J. H., Kieninger, M., Kloch, A., Pelan, S., Robertson, E., Pedersen, A. B., Abreu-Goodger, C., Buck, A. H., & Blaxter, M. (2023). Ancient diversity in host-parasite interaction genes in a model parasitic nematode. Nature Comm. 14, 7776.

Storer, J., Hubley, R., Rosen, J., Wheeler, T. J., & Smit, A. F. (2021). The Dfam community resource of transposable element families, sequence models, and genome annotations. Mobile DNA 12, 2.

Strome, S., Kelly, W. G., Ercan, S., & Lieb, J. D. (2014). Regulation of the X chromosomes in *Caenorhabditis elegans*. Cold Spring Harbor Perspectives in Biology 6, a018366.

Sudhaus, W., & Kiontke, K. (2007). Comparison of the cryptic nematode species *Caenorhabditis brenneri sp. n.* and *C. remanei* (Nematoda: Rhabditidae) with the stem species pattern of the *Caenorhabditis* Elegans group. Zootaxa 1456, 45–62.

Sun, S., Kanzaki, N., Dayi, M., Maeda, Y., Yoshida, A., Tanaka, R., & Kikuchi, T. (2022). The compact genome of *Caenorhabditis niphades n. sp*., isolated from a wood-boring weevil, *Niphades variegatus*. BMC Genomics 23, 765.

Ter-Hovhannisyan, V., Lomsadze, A., Chernoff, Y. O., & Borodovsky, M. (2008). Gene prediction in novel fungal genomes using an *ab initio* algorithm with unsupervised training. Genome Research 18, 1979–1990.

Teterina, A. A., Willis, J. H., Lukac, M., Jovelin, R., Cutter, A. D., & Phillips, P. C. (2023). Genomic diversity landscapes in outcrossing and selfing *Caenorhabditis* nematodes. PLoS Genetics 19, e1010879.

Thomas, J.H., Kelley, J.L., Robertson, H.M., Ly, K. & Swanson, W.J. (2005). Adaptive evolution in the SRZ chemoreceptor families of *Caenorhabditis elegans* and *Caenorhabditis briggsae*. Proceedings of the National Academy of Sciences 102, 4476–4481.

Thomas, J. H. (2006). Analysis of homologous gene clusters in *Caenorhabditis elegans* reveals striking regional cluster domains. Genetics 172, 127–143.

Toker, I.A., Ripoll-Sánchez, L., Geiger, L.T., Sussfeld, A., Saini, K.S., Beets, I., Vértes, P.E., Schafer, W.R., Ben-David, E. and Hobert, O., 2025. Divergence in neuronal signaling pathways despite conserved neuronal identity among *Caenorhabditis* species. Current Biology 35, 2927–2945.

True, J.R., & Haag, E.S. (2001) Developmental system drift and flexibility in evolutionary trajectories. Evol Dev. 3, 109–19.

Wang, J., Chen, P.-J., Wang, G. J., & Keller, L. (2010). Chromosome size differences may affect meiosis and genome size. Science 329, 293.

Wenke, T., Döbel, T., Sörensen, T. R., Junghans, H., Weisshaar, B., & Schmidt, T. (2011). Targeted identification of short interspersed nuclear element families shows their widespread existence and extreme heterogeneity in plant genomes. The Plant Cell 23, 3117–3128.

Woodruff, G. C., & Teterina, A. A. (2020). Degradation of the repetitive genomic landscape in a close relative of *Caenorhabditis elegans*. Molecular Biology and Evolution 37, 2549–2567.

Xiong, W., He, L., Lai, J., Dooner, H. K., & Du, C. (2014). HelitronScanner uncovers a large overlooked cache of Helitron transposons in many plant genomes. Proc Natl Acad Sci USA 111, 10263–10268.

Yang, Z. (1994). Maximum likelihood phylogenetic estimation from DNA sequences with variable rates over sites: approximate methods. Journal of Molecular Evolution 39, 306–314.

Yoshida, K., Rödelsperger, C., Röseler, W., Riebesell, M., Sun, S., Kikuchi, T., & Sommer, R. J. (2023). Chromosome fusions repatterned recombination rate and facilitated reproductive isolation during *Pristionchus* nematode speciation. Nature Ecology & Evolution 7, 424–439.

Zdraljevic, S., Walter-McNeill, L., Bruni, G. N., Bloom, J. S., Leighton, D. H. W., Collins, J. B., Marquez, H., Alexander, N., & Kruglyak, L. (2024). Divergent *C. elegans* toxin alleles are suppressed by distinct mechanisms. bioRxiv 10.1101/2024.04.26.591160.

Zhang, C., Rabiee, M., Sayyari, E., & Mirarab, S. (2018). ASTRAL-III: polynomial time species tree reconstruction from partially resolved gene trees. BMC Bioinformatics 19(Suppl 6), 153.

Zhao, Z., Boyle, T. J., Bao, Z., Murray, J. I., Mericle, B., & Waterston, R. H. (2008). Comparative analysis of embryonic cell lineage between *Caenorhabditis briggsae* and *Caenorhabditis elegans*. Developmental Biology 314, 93–99.

