## Supplementary Figures for "Conservative evolution of genetic and genomic features in *Caenorhabditis becei*, an experimentally tractable gonochoristic worm"

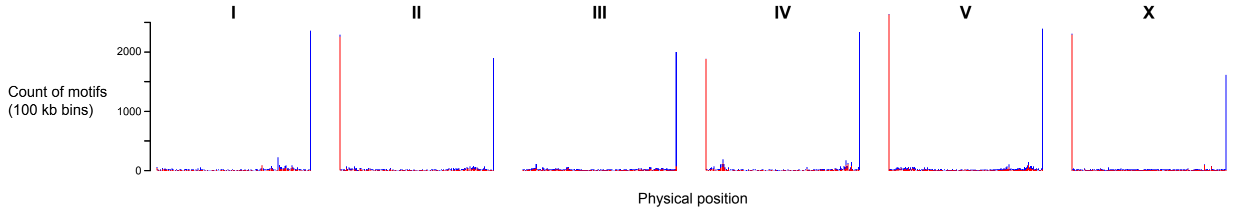


**Figure S1.** Chromosomes end with oriented telomere sequences in most cases. The plot shows stacked histograms of the counts of TTAGGC (blue) and GCCTAA (red), in 100kb bins along each chromosome. The left ends of chromosomes I and III lack telomere sequences.

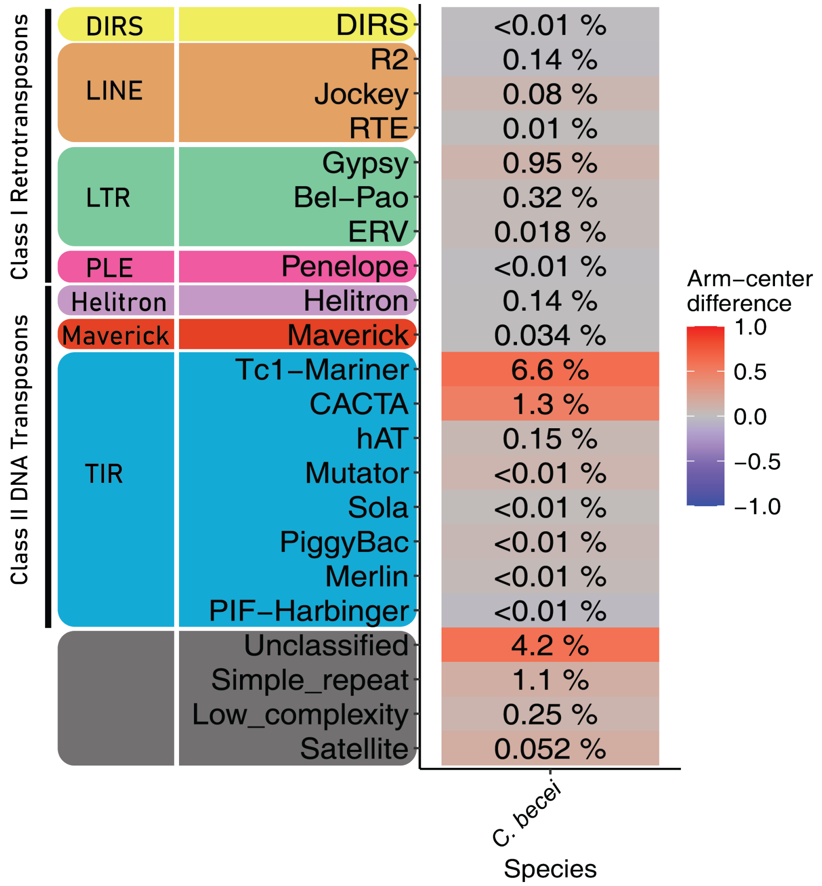


**Figure S2.** Transposable element superfamilies in *C. becei*. Each row represents a transposable element (TE) superfamily, grouped by higher-level taxonomic categories and colored by repeat order. Colored boxes next to each superfamily indicate the arm-center difference (Cohen's d), calculated as the difference in mean repeat density between chromosome arms (normalized position ≥ 0.25) and centers (normalized position < 0.25), divided by the pooled standard deviation. Positive values (red) reflect higher repeat density in chromosome arms, negative values (blue) indicate enrichment in chromosome centers, and values near zero are shown in grey. Numbers within boxes show the percentage of the genome occupied by each superfamily.

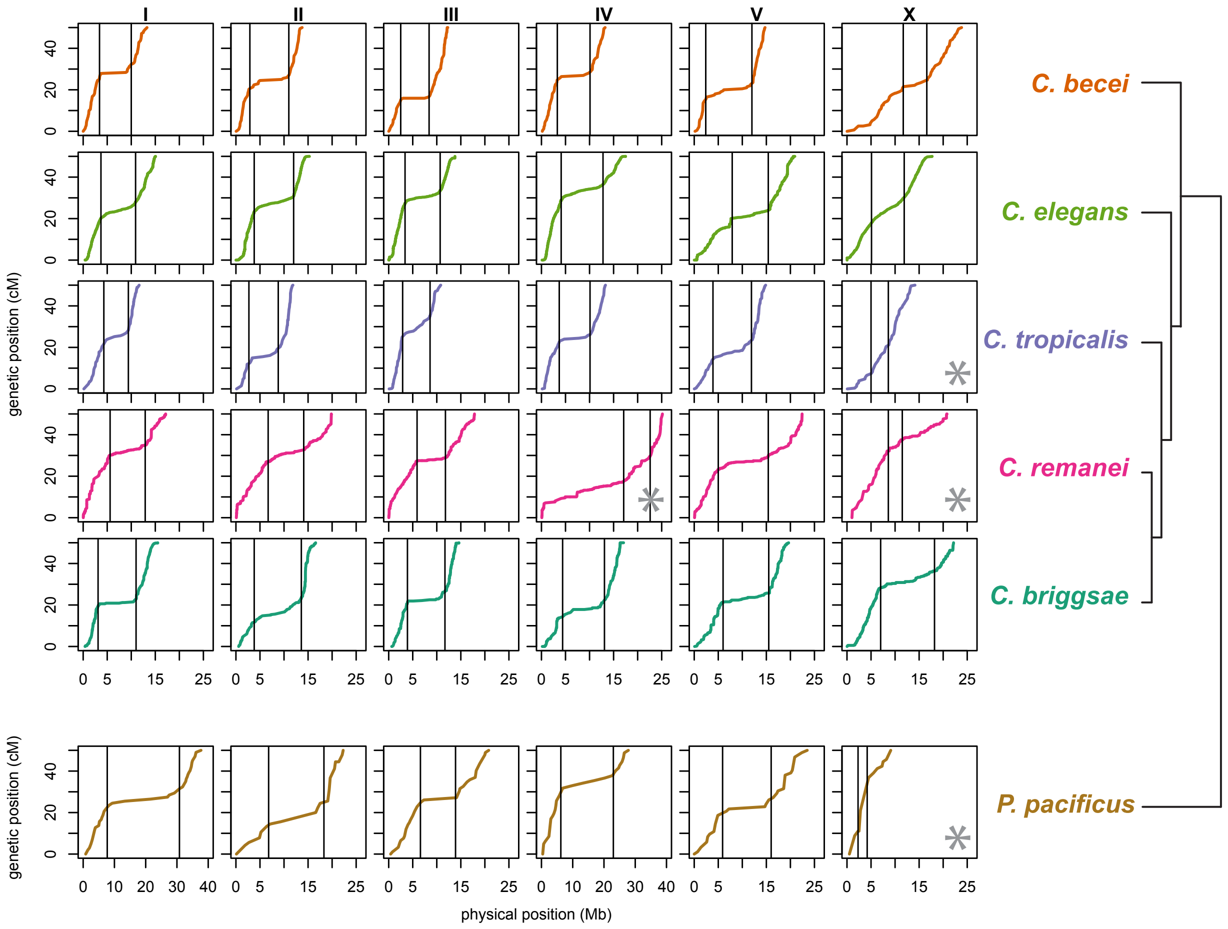


**Figure S3.** Domains in *C. becei* and other species. Marey maps for each of the chromosomes in six species with genetic map data. Vertical lines mark the estimated positions of chromosome domain boundaries. The x-axis is the physical position along the chromosome, in Mb, and the y-axis is the genetic position, in cM, after rescaling each map to 50 cM total length. The x-axis runs from 0 to 26 Mb in each plot, with the exceptions of *P. pacificus* chromosomes I and IV, which are much longer. As described in Table S2, four chromosome maps (marked with asterisks) do not have the expected domain structure: *C. tropicalis* X, *C. remanei* IV and X, and *P. pacificus* X.

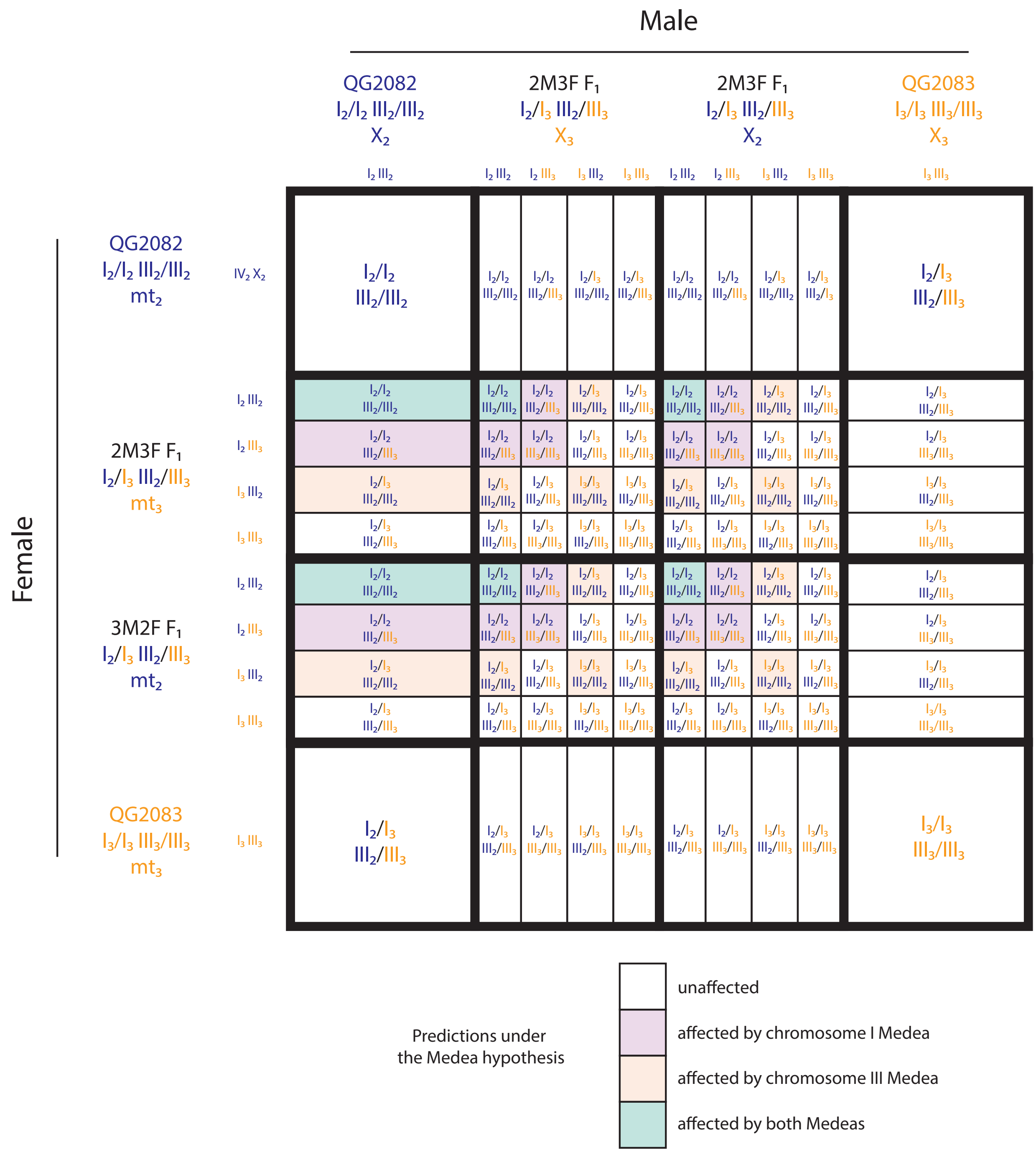


**Figure S4.** Sixteen Punnett squares showing the expected frequencies of affected progeny under a model of *Medea* elements on chromosomes I and III in QG2083 that independently affect QG2082-homozygous progeny of heterozygous mothers.

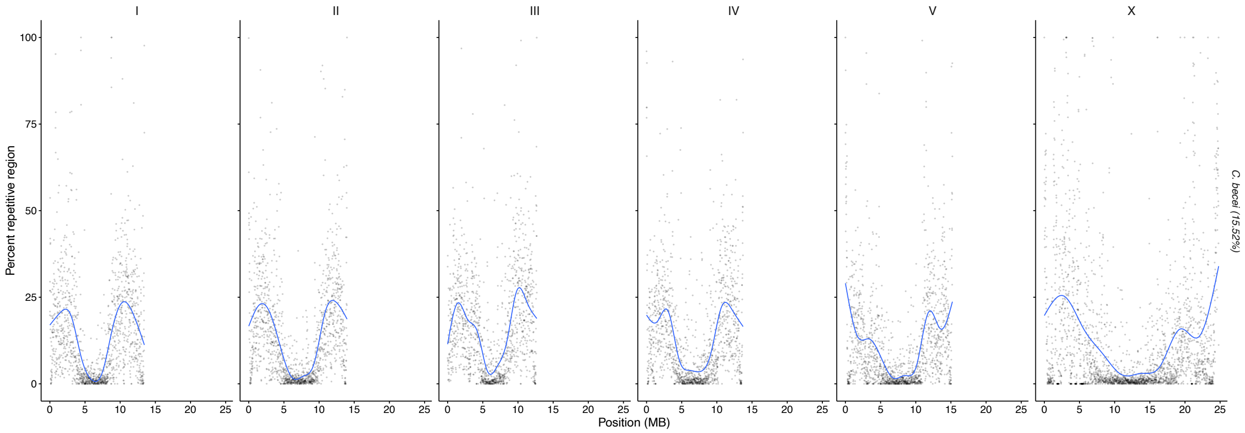


**Figure S5**. Global repetitive element landscape across 10-kb windows along the length of the chromosomes in *C. becei*. Blue lines show the smoothed trend in repeat density, generated by fitting a generalized additive model to the data.

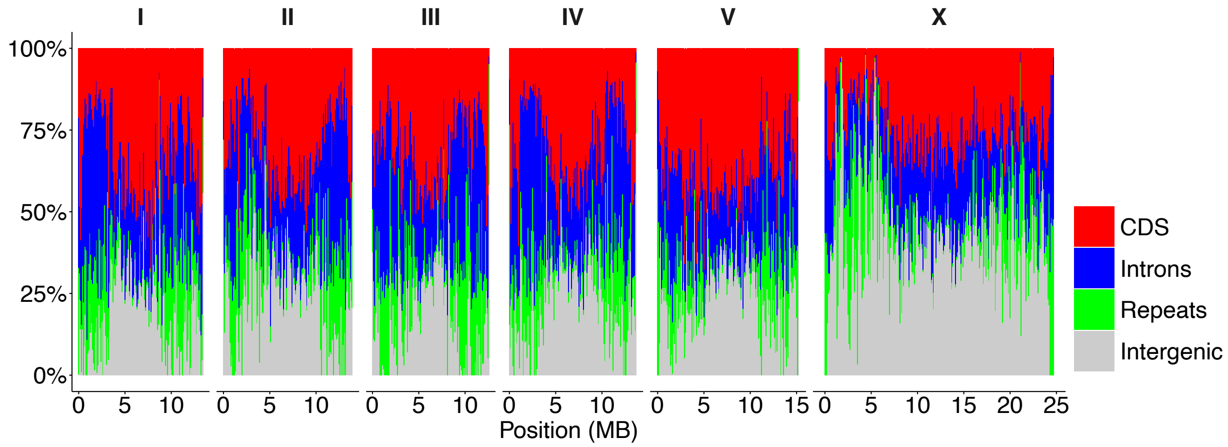


**Figure S6.** Genomic feature composition across the chromosomes of *C. becei***.** The proportion of each 100-kb window occupied by coding (CDS, red), intronic (blue), repetitive (green), and intergenic (gray) bases is shown along each chromosome. Bars are stacked to sum to 100% per window. Intergenic regions represent sequence not annotated as CDS, intron, or repeat.

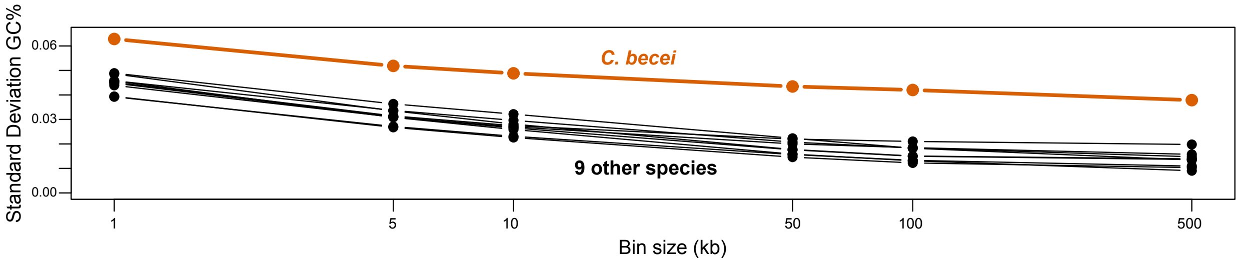


**Figure S7.** The *C. becei* genome is unusually variable in its local GC percentage. Here the standard deviation of GC% across the genome is shown as a function of the length scale over which GC is measured, in bins of 1 to 500 kb. The spacing along the x-axis is logged. The nine species shown in comparison to *C. becei* are those plotted in Figure 5.

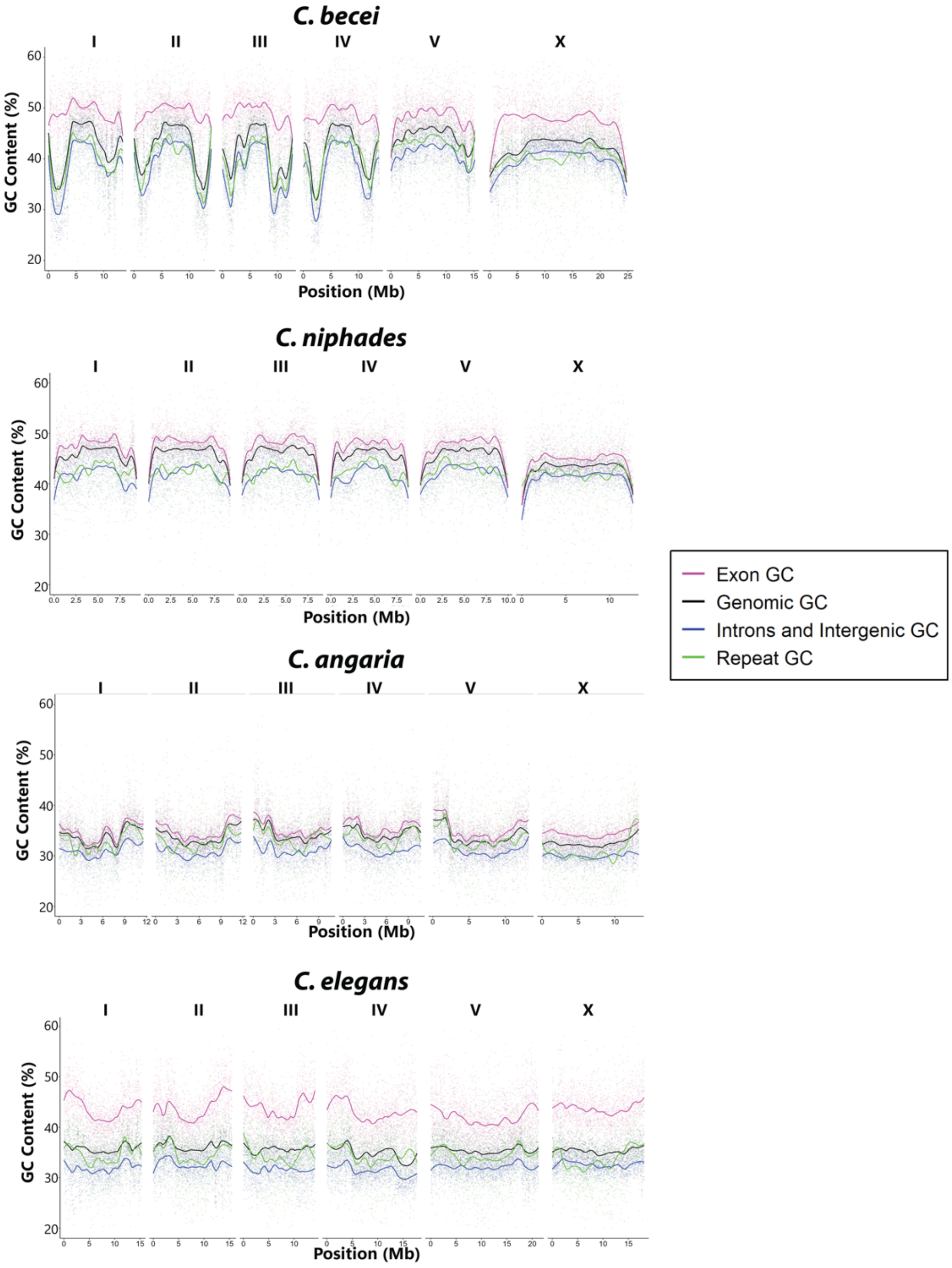


**Figure S8.** GC content of genomic features in *Caenorhabditis* species. GC content was calculated from the counts of G+C divided by the total number of bases of the feature within non-overlapping 10 kb windows along the length of the chromosome, with LOESS-fitted lines (span = 0.2). *C. elegans* here represents the relatively homogenous Elegans Group species.

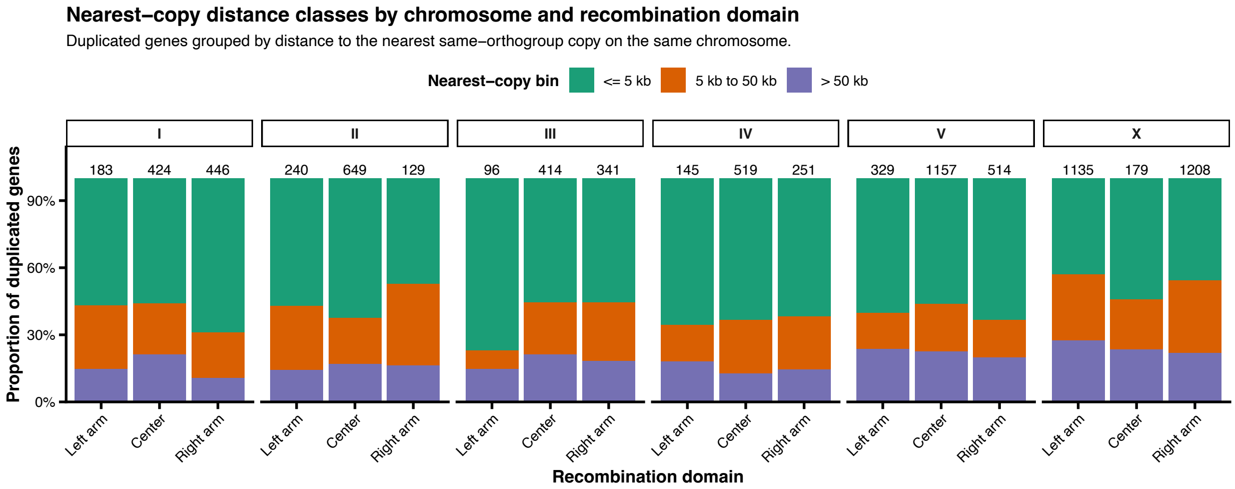


**Figure S9.** Distances from each multicopy gene to its nearest same-orthogroup neighbor.

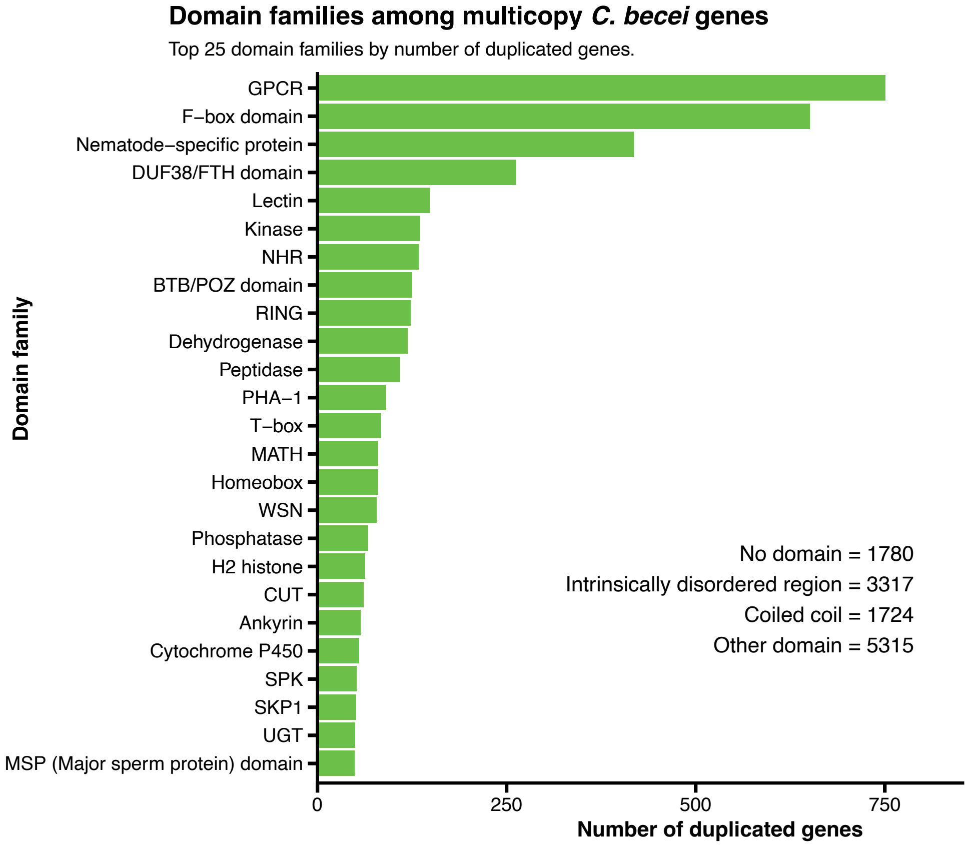


**Figure S10.** Domain families present in multicopy genes in *C. becei*. The histogram includes the 25 most prevalent curated domain families by gene count. Each bar shows the number of distinct multicopy genes containing that domain family; a gene is counted at most once within a given family but may contribute to more than one family if it contains multiple distinct domain families. Generic structural categories of intrinsically disordered regions and coiled coils, as well as genes lacking a retained domain annotation, are reported separately. “Other domain” represents the sum of gene-by-domain-family memberships across all curated domain families outside the ones shown individually; because a gene can contain more than one distinct domain family, this value does not represent a count of unique genes.

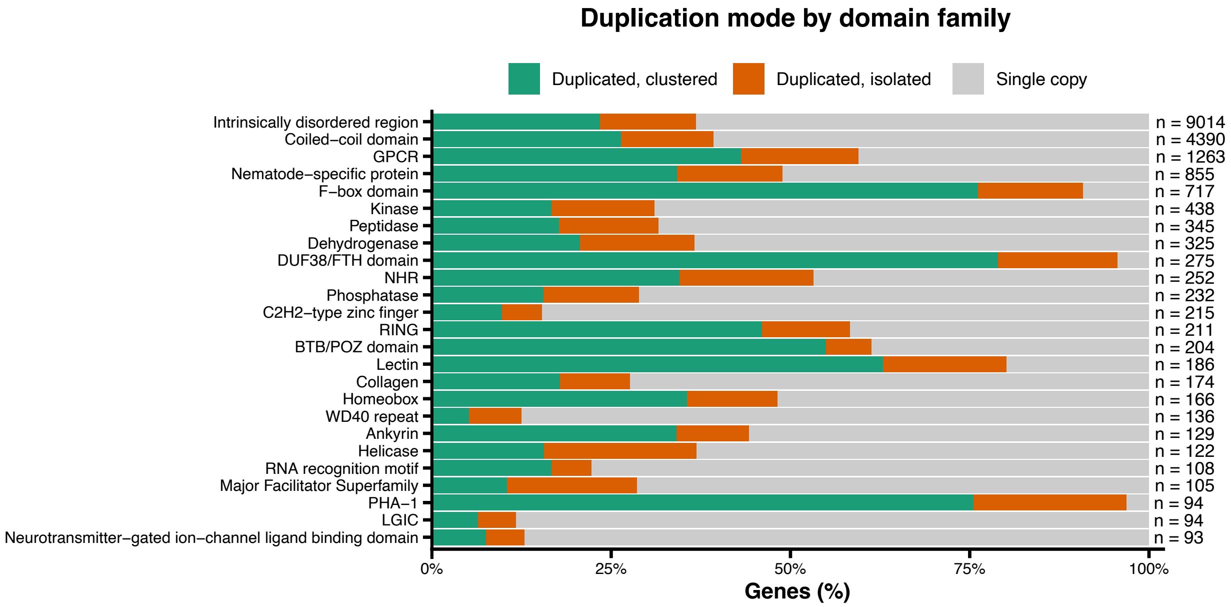


**Figure S11.** Domain families vary in their distribution across single-copy and multicopy genes. For the 25 most prevalent domain families, the bars show the proportion of genes that contain the domain and are single- or multicopy according to the Orthofinder classification. Multicopy genes are broken up into isolated and clustered counts, according to whether the gene's nearest same-orthogroup neighbor is nearby (less than 50 kb) or distant. The single-copy category includes small numbers of genes classified by Orthofinder as unassigned but that carry the specified domains.

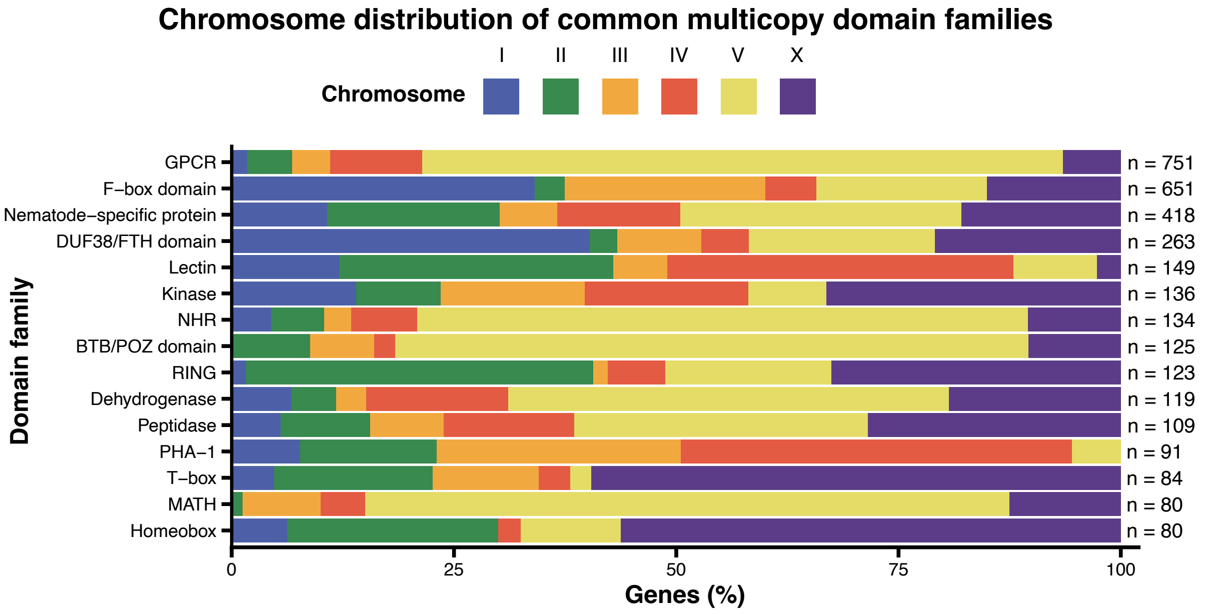


**Figure S12.** Domain families in multicopy genes vary in their distribution across chromosomes. For the 15 domains most represented among multicopy genes (excluding intrinsically disordered region and coiled coil domain), the bars show the proportion of genes that contain the domain on each chromosome (colored as in Figures 1 and 6). Multicopy genes containing GPCR, NHR, BTB/POZ, and MATH domains are particularly abundant on chromosome V, while T-box and Homeobox-containing multicopy genes are concentrated on the X chromosome.
